# Formamide denaturation of double-stranded DNA for fluorescence *in situ* hybridization (FISH) distorts nanoscale chromatin structure

**DOI:** 10.1101/2024.03.01.582995

**Authors:** Anne R. Shim, Jane Frederick, Emily M. Pujadas, Tiffany Kuo, I Chae Ye, Joshua A. Pritchard, Cody L. Dunton, Paola Carrillo Gonzalez, Nicolas Acosta, Surbhi Jain, Nicholas M. Anthony, Luay M. Almassalha, Igal Szleifer, Vadim Backman

## Abstract

As imaging techniques rapidly evolve to probe nanoscale genome organization at higher resolution, it is critical to consider how the reagents and procedures involved in sample preparation affect chromatin at the relevant length scales. Here, we investigate the effects of fluorescent labeling of DNA sequences within chromatin using the gold standard technique of three-dimensional fluorescence *in situ* hybridization (3D FISH). The chemical reagents involved in the 3D FISH protocol, specifically formamide, cause significant alterations to the sub-200 nm (sub-Mbp) chromatin structure. Alternatively, two labeling methods that do not rely on formamide denaturation, resolution after single-strand exonuclease resection (RASER)-FISH and clustered regularly interspaced short palindromic repeats (CRISPR)-Sirius, had minimal impact on the three-dimensional organization of chromatin. We present a polymer physics-based analysis of these protocols with guidelines for their interpretation when assessing chromatin structure using currently available techniques.

## Introduction

Chromatin organizes into hierarchical structures associated with maintaining cell-type-specific gene expression. At the nucleosomal scale (<1 Kbp [1]), epigenetic modifications control access for transcriptional proteins to genes of interest; at the level of topologically associating domains (TADs, 100 Kbp – 1 Mbp [2,3]), architectural proteins maintain proximity between linearly distant enhancers and promoters; and at the level of A/B compartments (1 – 100 Mbp [4,5]), the phase separation between gene-rich and gene-poor regions of chromatin may increase interactions between genes and proteins [6]. Recent work has established the existence of chromatin domains with sizes between the nucleosome and TAD length scales that are functionally important for regulating transcription [7–9]. Within chromatin packing domains, the chromatin polymer has a fractal-like behavior which is quantified by the power-law scaling parameter, *D* (*M* ∼ *r^D^*, where *M* is mass, quantified for chromatin by the number of base pairs, and *r* is the radius of the spherical volume under investigation) which is related to the contact scaling *s* observed in high-throughput conformation capture and oligo-paint measures of TADs [8,10,11]. Changes in chromatin scaling behavior have been correlated with altered gene expression patterns [12–14], phenotypic plasticity [14], carcinogenesis [15–17], chemotherapeutic efficacy [12,14], and reduced survival in cancer patients [14,18,19].

As a reflection of the considerable interest in understanding how aberrant genome organization leads to disease, there has been a massive expansion in imaging technologies and sample preparation protocols that allow chromatin to be visualized at super-resolution and smaller length scales [10,20–25]. The most common methods employ fluorescence *in situ* hybridization (FISH) techniques or CRISPR/dCas9-based labeling strategies to identify the locations of specific loci of interest. DNA (∼1 nm) and histones (∼10 nm) are well below the diffraction limit of light (∼200 nm) and therefore require labeling with fluorescent proteins or other large probes [10]. These specialized labeling techniques could perturb chromatin structure, which poses a paradoxical challenge: it is only with labels that a locus can be identified, but the chromatin structure of the labeled locus is not necessarily representative of its unlabeled, native state.

Prior work with both fluorescence and electron microscopy has demonstrated that the ultrastructure of the genome is sensitive to such labeling methods [26–28], but how these protocols relate to the structure of live cells is poorly understood. To address this knowledge gap, we investigate global nuclear chromatin structure before and after labeling with a gold-standard loci-specific method, 3D DNA FISH [29], to identify whether existing protocols are disrupting the native chromatin structure. Further, we probe which specific reagents and steps in the 3D FISH sample preparation protocol are deleterious to chromatin structure across the nucleus. We find that formamide exposure is the predominant cause of widespread alterations to chromatin domains. Finally, we evaluate other DNA-sequence labeling methods that do not require denaturation of double-stranded DNA, such as RASER-FISH [30] and CRISPR-Sirius [31], and show that these protocols are far less damaging to chromatin packing domains when compared to 3D FISH.

## Results

As chromatin packing domains have an average radius of ∼80 nm [11], the domain conformation within these subdiffractional structures cannot be effectively resolved by most conventional optical techniques. However, though the structure of individual domains cannot be resolved, it is possible to obtain information about their sub-diffractional organization in live cells by utilizing label-free nanoscopic sensing modalities such as Partial Wave Spectroscopic (PWS) microscopy [8,32]. In each diffraction-limited pixel, PWS microscopy captures the signal originating from the heterogeneity of the nanoscale molecular density distribution [32]. Due to its sensitivity to length scales between 20-200 nm (kin to sub-Mbp length scales) [33], the resulting signal is sensitive to changes in conformation within chromatin packing domains down to the size of the chromatin chain [11,21]. As a result, PWS microscopy offers several advantages over traditional super-resolution microscopy that are vital for this study: 1) it is label-free, allowing for careful study of nanoscale chromatin organization without perturbation; 2) both live and fixed cells can be imaged, which provides an opportunity to compare labeling protocols requiring fixation to the original, native chromatin structure; and 3) due to the fast acquisition time (<5 seconds) and a wide field of view (∼10,000 µm^2^), it is a high-throughput technique that provides increased statistical power by simultaneously imaging hundreds of cells within minutes [8,32].

Importantly, PWS microscopy is sensitive to length scales containing chromatin packing domains [32,33], which are spatially separable regions that follow power-law scaling behavior [8,11]. Analysis of PWS microscopy images allows for the identification of chromatin packing domains and the measurement of the statistics of the chromatin chain conformation within these domains [8,32]. In polymer physics, the space a polymer occupies (the radius of gyration, *r_g_*) can be related to the size of the polymer (the number of bond segments, *N*) through the Flory exponent, *ν*, which describes the scaling behavior (*r_g_* ∼ *N^v^*) [34]. As chromatin is a complex polymer, within a chromatin packing domain the mass of the chromatin contained within the domain *M* is related to the size of the domain *r* through the scaling exponent *D* (*M* ∼ *r^D^*). Chromatin packing domains were first identified in electron microscopy images by finding DNA-dense domain centers, and then analyzing where the domain stops following power-law scaling to determine the domain boundary [8,11]. Because the only factor that identifies a chromatin packing domain is the ability to follow power-law scaling, every packing domain can have a unique mass (*M*), radius (*r*), or scaling behavior (*D*) [8,11]. Given that *D* is the inverse of *ν*, in the absence of spatial constraints *D* can theoretically range from 5/3, if the chromatin within a packing domain exhibits an expanded coil state (self-avoiding walk) [35], to 3, for chromatin organized into a compact globule state [4,36]. However, polymers in a Θ solvent, such as chromatin, have the properties of a random walk (*D* ∼ 2) [37,38], therefore values below this are not a feasible model for chromatin organization within the nucleus [39,40], limiting the theoretical range to 2 < *D* < 3 [41,42]. Consistent with this finding, past studies quantified that the typical physiological range of *D* is 2.2-2.8, depending on the cell line [8,11,43].

### Formamide significantly alters chromatin structure in 3D FISH-labeled cells

Since PWS microscopy can visualize the nanoscale chromatin organization of whole nuclei in live and fixed cell populations, we imaged cells throughout each step in 3D FISH labeling protocols to determine their effects on nuclear structure. To confirm that the standard 3D DNA FISH protocol [29] is functional for the detection of specific genomic loci, we utilized this complete protocol for labeling chromosome 3. With this protocol, we were able to attain clear confocal images of foci within multiple nuclei (Fig 1A), which confirms that the protocol works as intended and that the altered chromatin domains are an unintended consequence of the 3D FISH labeling. Having demonstrated the suitability of this protocol for loci identification, we then performed mock staining using the typical 3D FISH protocol. The whole nuclear changes induced by each step were then compared to live untreated cells and cells fixed with 4% paraformaldehyde (PFA) for 10 minutes, which had previously been shown to maintain a correlation with the structures observed in live cells [44,45]. We conducted nuclear measurements comparing the effects of each step in two ways. First, we measured changes due to the cumulative effect of the protocol at each step (Fig 1B, light blue), to investigate whether any steps mitigate downstream changes within the protocol. Next, we investigated changes that occurred due to the individual reagents used in the 3D FISH protocol (Fig 1B, dark blue) by fixing the cells with PFA, performing any necessary wash steps, and then adding the reagent of interest.

**Fig 1.**
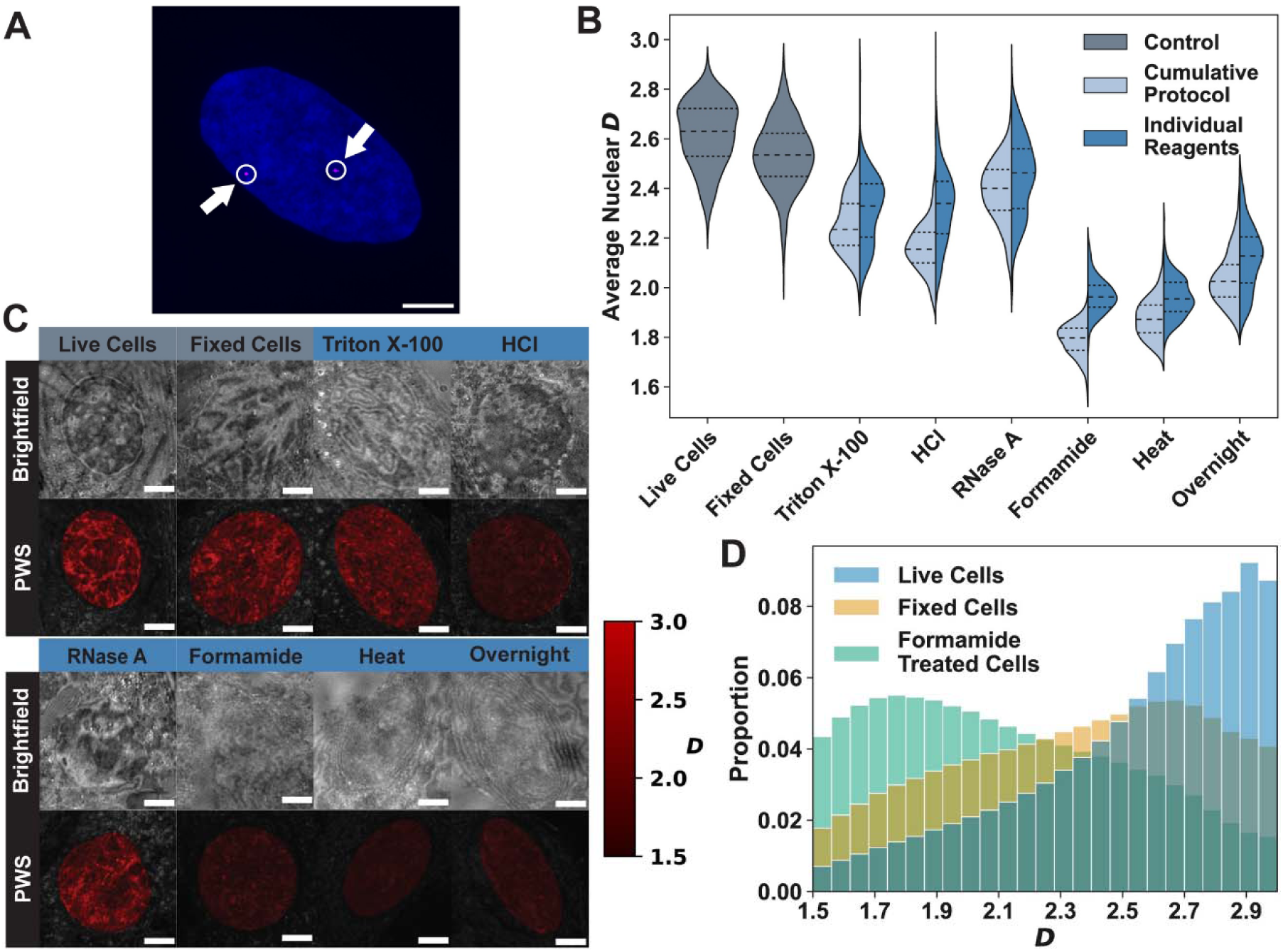
FISH damages chromatin domain structure across the nucleus. **A)** Representative image of a nucleus labeled with a chromosome 3 probe (magenta) and a DAPI nuclear counterstain (blue). The two chromosome 3 foci are denoted by a circle and an arrow. The image is a max projection of ∼15 planes within a z-stack. Scale bar: 5 µm. **B)** The average *D* for each nucleus undergoing the cumulative 3D FISH protocol (following the full protocol by performing each step consecutively; light blue) and treated with the individual 3D FISH reagents (fixing cells first, then performing the step listed; dark blue) compared to a live cell control and a 4% PFA fixed cell control (gray). The 3D FISH protocol has seven main steps: (1) cell fixation with 4% PFA, (2) permeabilization with Triton X-100, (3) deproteinization with hydrochloric acid (HCl), (4) degradation of RNA using RNase A, (5) lowering DNA melting point through formamide treatment, (6) heat denaturation of DNA, and (7) hybridization of the probe to DNA during overnight incubation. There are no protective benefits to performing the protocol in order. After completing the whole protocol, the average *D* of nuclei is far below control cells. All data is an average of between 240-800 nuclei, across three biological replicates. Dashed lines within violins denote the 75^th^ percentile, median, and 25^th^ percentile from top to bottom. **C)** A comparison of brightfield and PWS microscopy images (brighter red indicates higher *D*) after treatment with each reagent. Some treatments of the 3D FISH protocol cause nuclei to appear vastly different, even on brightfield images. Since the nuclear membrane is visible after heat treatment and overnight incubation, it is unlikely that these treatments are completely degrading the nuclear membrane. However, it is impossible to visually detect the location of the nucleus after formamide treatment without using a nuclear counterstain. Scale bar: 5 µm. **D)** The probability distribution function for *D* of each pixel within all analyzed nuclei in the live cell control, the fixed cell control, and the formamide-treated cells. Formamide causes the PDF to shift towards a random distribution, centered around *D* ≈ 2.1. Formamide lowers the most probable *D* to ∼1.8, much lower than the values observed in healthy cells.

Although past studies showed that there was a change in chromatin structure only after heat denaturation [30,26,27,46], we found that all phases of the 3D FISH protocol cause significant changes to nanoscale chromatin structure in comparison to living cells (Figs 1B-C and Table 1). Even brightfield imaging shows that nuclei are visibly damaged at every step after Triton X-100 treatment (Fig 1C), without any evidence of mitigating effects on downstream perturbations in the full sequence of sample preparation (Fig 1B). Indeed, the cumulative effects compound to a decrease in *D* when compared to the individual reagents in every step (Fig 1B and Table 1), indicating that rather than having a protective effect, the full 3D FISH protocol results in a cumulative loss of information in comparison to the live-cell state.

**Table 1.** Statistics of the change in D resultant from each 3D FISH reagent.

| Reagent/<br>Treatment | Average Nuclear $D \pm$<br>Standard Deviation | | % Change from Live<br>Cells | | % Change from Fixed<br>Cells | | % Difference<br>Between<br>Cumulative<br>Protocol and<br>Individual<br>Reagents |
| --- | --- | --- | --- | --- | --- | --- | --- |
|  | Cumulative<br>Protocol | Individual<br>Reagent | Cumulative<br>Protocol | Individual<br>Reagent | Cumulative<br>Protocol | Individual<br>Reagent |  |
| **Live Cells** | 2.62±0.13 (N=801 nuclei) | | N/A | | N/A | | N/A |
| **10 min**<br>**PFA** | 2.53±0.13 (N=554 nuclei) | | -3.23% (p=7.82x10<sup>-28</sup>) | | N/A | | N/A |
| **10 min**<br>**Triton X-100** | 2.25±0.11<br>(N=285<br>nuclei) | 2.32±0.15<br>(N=297<br>nuclei) | -13.97%<br>(p=4.01x<br>10<sup>-227</sup>) | -11.43%<br>(p=1.62x<br>10<sup>-160</sup>) | -11.10%<br>(p=1.66x<br>10<sup>-135</sup>) | -8.47%<br>(p=1.68x<br>10<sup>-80</sup>) | 2.92%<br>(p=2.21x10<sup>-7</sup>) |
| **5 min**<br>**HCl** | 2.16±0.10<br>(N=318<br>nuclei) | 2.33±0.15<br>(N=280<br>nuclei) | -17.46%<br>(p<10<sup>-308</sup>) | -11.14%<br>(p=1.28x<br>10<sup>-148</sup>) | -14.71%<br>(p=2.28x<br>10<sup>-217</sup>) | -8.18%<br>(p=6.96x<br>10<sup>-73</sup>) | 7.37%<br>(p=5.69x10<sup>-46</sup>) |
| **45 min**<br>**RNase A** | 2.39±0.13<br>(N=314<br>nuclei) | 2.45±0.16<br>(N=307<br>nuclei) | -8.80%<br>(p=8.74x<br>10<sup>-120</sup>) | -6.54%<br>(p=2.22x<br>10<sup>-64</sup>) | -5.75%<br>(p=1.47x<br>10<sup>-47</sup>) | -3.42%<br>(p=3.89x<br>10<sup>-15</sup>) | 2.45%<br>(p=2.79x10<sup>-5</sup>) |
| **1 hour**<br>**Formamide** | 1.79±0.06<br>(N=314<br>nuclei) | 1.97±0.07<br>(N=241<br>nuclei) | -31.63%<br>(p<10<sup>-308</sup>) | -24.90%<br>(p<10<sup>-308</sup>) | -29.35%<br>(p<10<sup>-308</sup>) | -22.39%<br>(p=1.98x<br>10<sup>-306</sup>) | 9.39%<br>(p=1.61x<br>10<sup>-118</sup>) |
| **3 min**<br>**Heat with**<br>**Hybridization**<br>**Buffer** | 1.88±0.08<br>(N=328<br>nuclei) | 1.96±0.08<br>(N=362<br>nuclei) | -28.36%<br>($p < 10^{-308}$) | -25.09%<br>($p < 10^{-308}$) | -25.97%<br>($p < 10^{-308}$) | -22.59%<br>($p < 10^{-308}$) | 4.46%<br>($p = 4.11 \times 10^{-36}$) |
| **12-15 hours**<br>**Overnight**<br>**Incubation** | 2.04±0.11<br>(N=316<br>nuclei) | 2.12±0.13<br>(N=240<br>nuclei) | -22.18%<br>($p < 10^{-308}$) | -19.08%<br>($p = 8.36 \times 10^{-289}$) | -19.59%<br>($p = 8.90 \times 10^{-291}$) | -16.38%<br>($p = 2.04 \times 10^{-193}$) | 3.91%<br>($p = 7.43 \times 10^{-13}$) |

Because of the large amount of damage observed after conducting the 3D FISH protocol, we sought to determine the point at which these changes were initiated. Therefore, we measured the change in *D* following each step in the protocol (Fig 1B and Table 1). First, we found that fixation alone causes a decrease in *D* (Δ*D* ≈ –0.08) which is smaller than the natural standard deviation of *D* in the live cell population (σ*_D_* ≈ 0.13) and the standard deviation within the fixed cells (σ*_D_* ≈ 0.13, Fig 1B and Table 1). All the steps after fixation caused a compression of the violin plots (σ*_D_* < 0.13, Fig 1B and Table 1, Cumulative Protocol), indicating that the population of cells undergoing mock preparation has less variability than would occur within the live-cell population. The largest changes in average chromatin scaling were due to permeabilization by Triton X-100 (live cell Δ*D* ≈ –0.37 and fixed cell Δ*D* ≈ –0.28) and deproteinization with hydrochloric acid (HCl) treatment (live cell Δ*D* ≈ –0.46 and fixed cell Δ*D* ≈ –0.37; Fig 1B and Table 1, Cumulative Protocol). On average, the lower *D* induced by both Triton X-100 and HCl treatments indicate a shift towards more loosely packed chromatin. This could be caused by either the increased flux of constituents between the nucleus and cytoplasm, which is facilitated by the permeabilized nuclear membrane and the reduced intracellular protein concentration, or molecular denaturation that can be associated with detergent/acid exposures. The RNA degradation step slightly increased *D* compared to the previous steps (live cell Δ*D* ≈ –0.23 and fixed cell Δ*D* ≈ –0.15; Fig 1B and Table 1, Cumulative Protocol) but this change was not sustained on further steps within the protocol.

While reagents used at the beginning of the protocol such as Triton X-100 and HCl caused widespread changes in *D*, we observed that formamide exposure was the most deleterious to the underlying nanoscale chromatin structure. In brightfield imaging, nuclei cannot be visually detected after formamide treatment. To understand what changes are occurring within nuclei, we compared the probability distribution function (PDF) of *D* (within each of the 250-nm voxels in the nuclei) to understand how local scaling is altered due to the most damaging step in the protocol (Fig 1D). Formamide not only caused the largest decrease in the average *D* (live cell Δ*D* ≈ –0.83 and fixed cell Δ*D* ≈ –0.74; Fig 1B and Table 1, Cumulative Protocol) but was also associated with a complete shift in the PDF of *D* values within the population, causing the most probable configuration within nuclei to decrease to ∼1.74 as compared to ∼2.88 for live cells (Fig 1D). For context, this is much lower than has previously been observed for viable cells [8,11] and would be consistent with a polymer in an expanded coiled state [47]. Although formamide resulted in disruption of nuclear structure, the nuclear membrane was visible at later steps of the protocol indicating that formamide does not degrade the nuclear membrane but interacts with the genetic material within the nucleus to change the polymeric structure of the genome.

Formamide is utilized within most FISH protocols because it (1) lowers the melting temperature of DNA and (2) stabilizes the transition from the active helical state of DNA to the denatured coil phase [48]. Mechanistically, formamide achieves this state as it forms stronger hydrogen bonds with water than other water-water or water-DNA hydrogen bonds, and therefore displaces DNA hydrate, forms hydrogen-bonded networks, and destabilizes the DNA helix [49]. However, the remaining denatured coil phase of DNA is structurally distinct from the physiological states of DNA within the nucleus and is much more homogenously folded. In the context of chromatin forming domains to achieve preferentially accessible structures [39], these findings indicate that formamide exposure results in homogeneous chromatin configurations that are not representative of the structures observed in live cells [8,11].

Next, we evaluated the effect of heat denaturation on chromatin organization following formamide exposure to see if this had any further effect on chromatin organization and notably did not see substantial changes in chromatin packing. It is also important to note that every step following formamide incubation continues to use formamide in its buffers, including heat denaturation and overnight incubation. While heat-mediated DNA-denaturation is not an inert process, the structure of chromatin was altered so significantly by the initial formamide treatment that heat denaturation did not have any significant reversal nor deleterious effects as measured by *D* (live cell Δ*D* ≈ –0.74 and fixed cell Δ*D* ≈ –0.66; Fig 1B and Table 1, Cumulative Protocol). Therefore, in every step of the 3D FISH protocol following formamide incubation, the DNA is likely to be in a stabilized coil state due to the strong formamide-water hydrogen bonds and locally folded at random compared with the active helical state. Overnight incubation modestly increased *D* (live cell Δ*D* ≈ –0.58 and fixed cell Δ*D* ≈ –0.50), however, the final state was consistent with a random walk polymer (*D* ≈ 2.04; Fig 1B and Table 1, Cumulative Protocol). In sum, the 3D FISH-labeled cells experienced a shift toward lower *D* values such that cells had an average nuclear *D* value of 2.04, as compared to live cells, which had an average *D* of 2.62.

As chromatin is a biopolymer that organizes following polymer scaling properties, this suggests that chromatin following 3D FISH preparation has the average properties of a random walk polymer (ν ∼ 1/*D* ∼ ½) [35]. Physiologically, the lowest *D* value observed in a packing domain within a nucleus is ∼2.2, as observed by electron microscopy [11]. This indicates that the 3D FISH protocol causes reorganization of chromatin such that it behaves in a non-physiological manner. In the context of genomic structures in the domain range (e.g. 100 Kbp – 1 Mbp domain), with the chromatin polymer being composed of monomers that are a single nucleosome each, the difference in observed domain radius from a *D* of ∼2.6 to ∼2 can be approximated using the relationship of *r_g_* ∼ *L*^1/*D*^. Here, *L* is the segment length, or the number of nucleosomes multiplied by the nucleosome diameter (∼11 nm), with ∼200 bp per nucleosome [1]. With this approximation, a 1.2 Mbp domain in live cells would occupy a volume with *r_g_ ∼* 72 nm, but upon preparation with 3D FISH, would be observed to have *r_g_ ∼* 256 nm for *D* = 2 and *r_g_ ∼* 456 nm for *D* = 1.8. These sizes are concordant with the findings from multiplexed DNA paint to probe the organization of topologically associated domains with 3D-FISH while using formamide denaturation [10,20].

### 3D FISH protocol optimization is unable to prevent formamide-induced damage to chromatin

Due to the large amount of damage observed after conducting the 3D FISH protocol, we sought to determine if modifications at various stages in sample preparation could offset these effects. FISH protocols are often optimized according to the cell type and the choice of probe to ensure high labeling efficiency and minimal impact on cellular structures. Therefore, we attempted to systematically adjust the steps to prevent formamide-induced damage or restore the structure after treatment. In principle, the greatest effect in structural preservation results during the process of fixation, as stronger fixatives or longer incubation times with fixatives can result in a more highly crosslinked or condensed chromatin structure that can withstand alteration via small molecules such as formamide. We performed measurements to determine the impact of fixation in the following manner: we (1) treated cells with the most-used fixative reagents/solutions and incubation times, (2) imaged the fixed cell populations with PWS, (3) treated all the fixed cells with the same formamide incubation treatment (2X SSC/50% formamide/0.1% Tween 20 for 30 minutes at room temperature), and (4) imaged the cells again with PWS. An ideal fixation step would induce minor changes in *D* inherently, reduce the change that is caused by formamide, and allow for a high labeling efficiency to facilitate fluorescence imaging of foci. Most 3D DNA FISH protocols use 4% PFA with short incubation times to prevent over-fixation through excessive crosslinking [50]. We evaluated incubation lengths of 10, 30, and 90 minutes with 4% PFA and found that increasing the incubation time reduced the change in *D* induced by fixation and by formamide treatment (Figs 2A-B and S1 Table).

**Fig 2.**
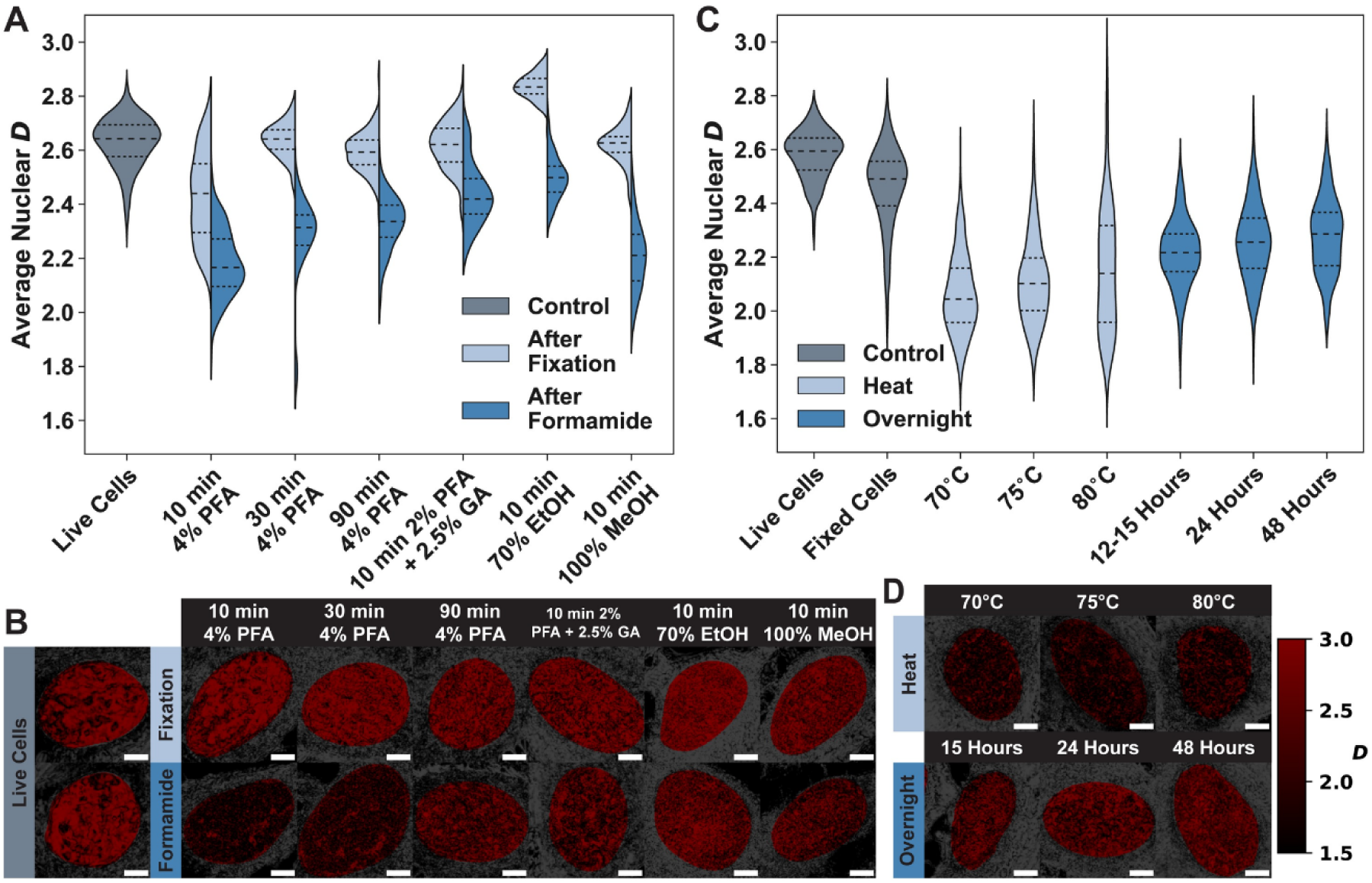
Optimization of the 3D FISH protocol does not prevent or reverse the formamide-induced chromatin structural alterations. **A)** Formamide causes changes to chromatin structure regardless of the mechanism or length of fixation. The most common fixative for FISH, 4% paraformaldehyde (PFA), is normally varied in the length of incubation with *in vitro* cell cultures to ensure that the cell remains stable over time and allows proper labeling of DNA with the desired probe. Increasing the incubation time indeed better preserves the original chromatin structure by reducing the change in *D* due to fixation. Other fixation reagents such as glutaraldehyde (GA) mixed with PFA, ethanol, and methanol seem to perform as well or better than increasing the incubation time. However, it is notable that ethanol fixation increases *D*, leading to formamide appearing to have less of an impact. Methanol, while having negligible effect during fixation, experiences the largest change because of formamide treatment. The 2% PFA with 2.5% GA solution has the least impact on *D* during fixation and does not experience as great a change from formamide. All data is an average of between 100-1000 nuclei, across three biological replicates. Dashed lines within violins denote the 75^th^ percentile, median, and 25^th^ percentile from top to bottom. **B)** Representative PWS microscopy images for live cell controls and fixed cells before and after formamide incubation (brighter red indicates higher *D*). Although the 10-minute 4% PFA fixation causes the largest change in average nuclear *D* values, qualitatively, the nuclei look the most like the chromatin structures visible in live cells. Most other stronger fixatives contain smaller high *D* clumps (chromatin packing domains) as compared to the live cells. After formamide treatment, however, none of the cells bear any resemblance to the original chromatin organization seen in live cells. Scale bar: 5 µm. **C)** The average nuclear *D* for two steps in the 3D FISH protocol that are commonly optimized to mitigate nuclear changes: heat denaturation and overnight incubation. Regardless of the temperature chosen for the heating step, *D* is close to 2, indicating that the temperature used for denaturing chromatin neither causes different chromatin structures nor reorganizes chromatin more than formamide treatment alone. Although the overnight incubation step shows a slight increase in *D* as the incubation time is increased, even a 48-hour incubation is not able to recover the original chromatin structure. Plotted data are from between 210 and 430 nuclei across three biological replicates. Dashed lines within violins denote the 75^th^ percentile, median, and 25^th^ percentile from top to bottom. **D)** PWS microscopy images were collected from nuclei treated with different temperatures or incubation times for the heat denaturation step and the overnight incubation step, respectively (brighter red indicates higher *D*). Scale bar: 5 µm.

Although PFA fixation is commonly used in FISH labeling, other solutions have been used to preserve intracellular structures. We assessed three additional solutions with reagents known to have a stronger impact on cells and which therefore could prevent damage to chromatin packing domains. First, we evaluated 10-minute incubation with a combination of 2% PFA and 2.5% glutaraldehyde (GA). Mixing GA with PFA is often done to increase sample stability and prevent DNA degradation over time [51–53]. We also evaluated two alcohol-based fixatives, 70% ethanol, and 100% methanol, which are common alternatives to aldehydes as they are thought to better preserve DNA structures [54] and could have higher labeling efficiency compared to aldehydes due to the lack of crosslinks. Of these fixative solutions, the 2% PFA and 2.5% GA solution had the most optimal results: the fixation itself caused almost no change to average nuclear *D* (live cell Δ*D* ≈ –0.01, n.s.) and was the least affected by formamide treatment (live cell Δ*D* ≈ –0.19 and fixed cell Δ*D* ≈ –0.18; Fig 2A and S1 Table). Ethanol fixation seemed to have the smallest change induced by formamide (live cell Δ*D* ≈ –0.13), however, this is due to the ethanol itself causing an increase in *D* (live cell Δ*D* ≈ 0.21) after fixation, resulting in a fixed cell Δ*D* ≈ –0.34 (Fig 2A and S1 Table). Lastly, methanol fixation had a negligible impact on *D* (live cell Δ*D* ≈ –0.01) but resulted in one of the largest changes in *D* after formamide treatment (live cell Δ*D* ≈ –0.42 and fixed cell Δ*D* ≈ –0.41; Fig 2A and S1 Table). The suboptimal results from alcohol-based fixative solutions are likely due to the fixation mechanism: ethanol and methanol both rely on cellular dehydration and condensation of proteins [51–53] which could leave DNA in alcohol-fixed cells susceptible to formamide. Although some fixative solutions mitigated the impact of formamide on chromatin structure, our results demonstrate that no reagent can completely prevent damage from formamide. The resulting chromatin was visually quite distinct from live cells (Fig 2B) and all of the fixed cells experienced a decrease in *D* of between 7-16% due to formamide (S1 Table).

After attempting to prevent damage through fixation, we hypothesized that heat denaturation or overnight incubation for hybridization could reduce the impact of the 3D FISH protocol on chromatin organization. As the chromatin structure was significantly altered by formamide treatment in all tested fixative solutions, we next investigated the effect of varying temperature during the heating step and varying incubation length for the overnight hybridization step on cells fixed with 4% PFA for 10 minutes. We incubated cells with the formamide denaturation treatment (2X SSC/50% formamide/0.1% Tween 20 for 30 minutes at room temperature), then performed the heat treatment in a hybridization buffer. We changed the temperature used for the heat denaturation step from 70-80°C (Figs 2C-D), as this is the common range of temperatures recommended for 3D FISH, and because the amount of formamide has been shown to lower the melting temperature of DNA linearly [55]. Each of the cell populations after heat treatment had an average population *D* of ∼2-2.1, showing a slight inverse relationship between the magnitude of change in *D* and the temperature. 70°C had the largest decrease (live cell Δ*D* ≈ –0.52 and fixed cell Δ*D* ≈ –0.40), followed by 75°C (live cell Δ*D* ≈ –0.47 and fixed cell Δ*D* ≈ –0.35), and 80°C (live cell Δ*D* ≈ –0.42 and fixed cell Δ*D* ≈ –0.30) (Fig 2C and S2 Table). Although increasing the temperature has a slight impact on the change induced by formamide, the benefit is minimal.

To evaluate whether damage to chromatin packing domains can be reversed by changing the overnight incubation step, we incubated cells with the formamide denaturation solution, performed heat denaturation at 75°C, moved the denatured cells to a humid chamber set at 37°C for hybridization, and measured nuclear chromatin organization after overnight (12-15 hours), 24-hour, or 48-hour incubation (Figs 2C-D). Increasing the incubation time led to more recovery of chromatin structure: the overnight incubation had the biggest decrease in *D* (live cell Δ*D* ≈ –0.37 and fixed cell Δ*D* ≈ –0.25) whereas the 24-hour (live cell Δ*D* ≈ –0.33 and fixed cell Δ*D* ≈ –0.20) and 48-hour (live cell Δ*D* ≈ –0.30 and fixed cell Δ*D* ≈ –0.18) incubations saw a marginally smaller change. However, the average nuclear *D* value only increased to ∼2.2 even with the longest incubation (Fig 2C and S2 Table). Additionally, although the average nuclear *D* value is within the physiologically relevant range, the chromatin packing domains in nuclei after overnight incubation were visually much smaller than those in live cells (Figs 2B and D). Taken together, our results indicate that modifying the denaturation and hybridization steps by changing temperature or incubation time can reverse some of the effects of formamide on chromatin packing domains, but critically, these modifications cannot completely recover the native structures seen in live cells.

### Alternate labeling methods that do not require formamide have minimal impact on chromatin organization

As optimization of the 3D FISH protocol could not preserve the native chromatin structure, we next applied other DNA labeling protocols that are either free of or use a minimal amount of formamide. We identified one fixed-cell FISH method, which has the benefit of using the same types of probes used in 3D FISH, and one live-cell CRISPR method, which allows for visualization of the motion of loci in live cells, albeit with lower resolution. For the non-denaturing FISH protocol, we selected RASER-FISH as it eliminates the need for formamide by creating ssDNA through exonuclease III digestion of UV-induced nicks in DNA [30,56] and has previously been used to compare TAD localization with DNA density based structural chromatin domains [7]. While there is a myriad of CRISPR-based labeling methods, we chose to utilize CRISPR-Sirius due to its enhanced guide RNA stability and brightness compared to earlier iterations of CRISPR labeling [31]. We performed PWS microscopy imaging on cells that were labeled with 3D FISH, RASER-FISH, and CRISPR-Sirius and compared resulting *D* values to those of both live cell and fixed cell controls to determine how global nuclear structure is impacted by each protocol (Figs 3A-B and Table 2). Labeling cells with a chromosome 3 probe using the 3D FISH protocol caused significant changes to chromatin (live cell Δ*D* ≈ –0.51 and fixed cell Δ*D* ≈ –0.49), whereas neither the RASER-FISH protocol with a chromosome 19 probe (live cell Δ*D* ≈ –0.03 and fixed cell Δ*D* ≈ –0.005) nor the CRISPR-Sirius method targeting XXYLT1 with a MS2 aptamer (live cell Δ*D* ≈ –0.08 and fixed cell Δ*D* ≈ –0.06) caused as much of a change (Fig 3A and Table 2). Notably, both RASER-FISH and CRISPR-Sirius had a similar effect as the fixation alone (live cell Δ*D* ≈ –0.02; Fig 3A and Table 2). This result indicates that at the population level, both techniques are better at preserving chromatin structure compared to 3D FISH.

**Fig 3.**
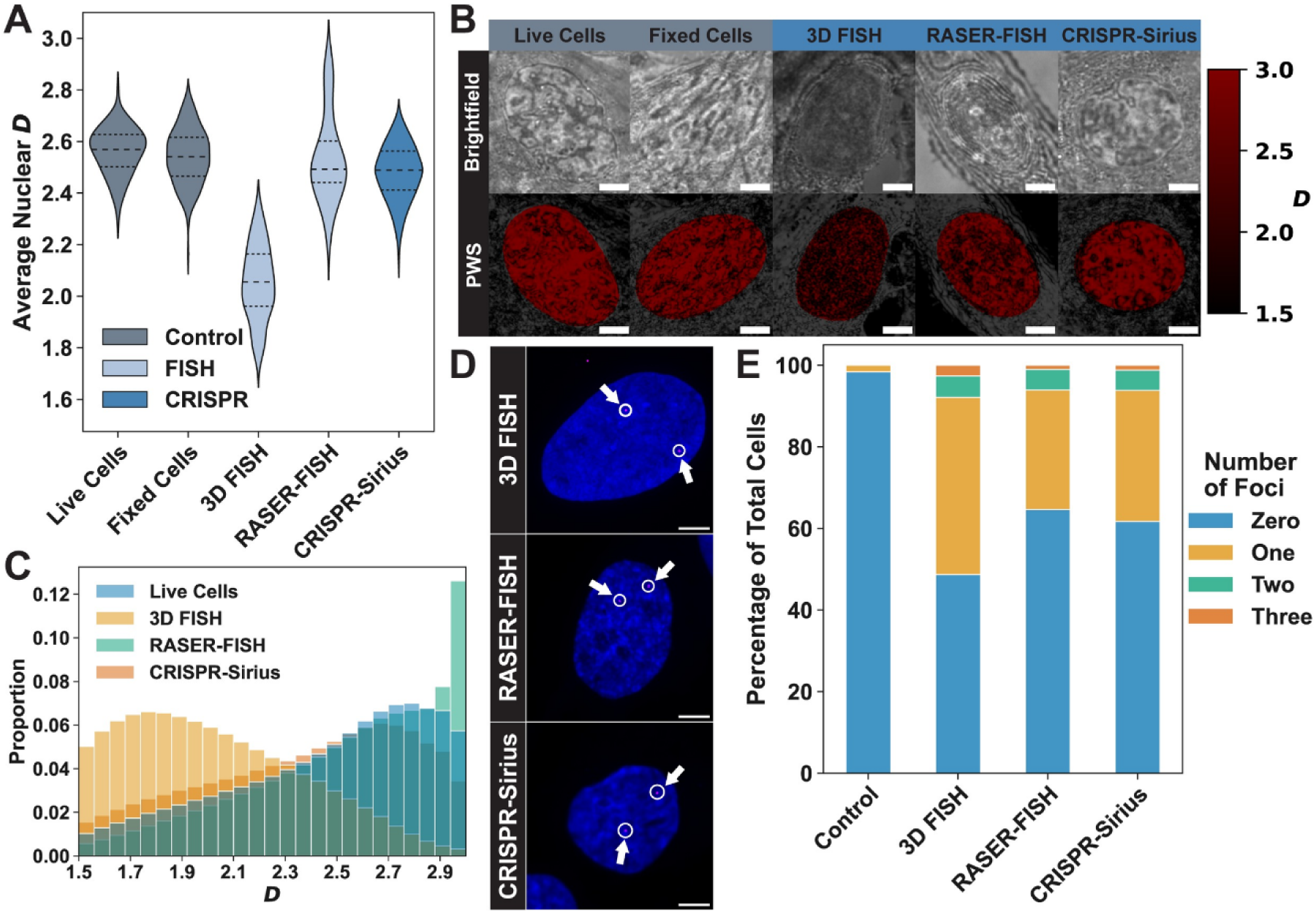
Alternate DNA sequence labeling methods have minimal impact on global nuclear chromatin organization. **A)** The average *D* for nuclei labeled with either 3D FISH (light blue), RASER-FISH (light blue), or CRISPR-Sirius (dark blue) compared to a live cell control (gray) and a 4% PFA fixed cell control (gray). All data is an average of between 70 and 170 nuclei, across three biological replicates. Dashed lines within violins denote the 75^th^ percentile, median, and 25^th^ percentile from top to bottom. **B)** A comparison of brightfield and PWS microscopy images (brighter red indicates a higher *D*) for each DNA labeling method. RASER-FISH labeled cells are visually different in both Brightfield and PWS, potentially due to the two fixation steps and the presence of formamide in the hybridization buffer. CRISPR-Sirius labeled cells look identical to the unstained live cells. Scale bar: 5 µm. **C)** The probability distribution function for *D* of each pixel within the analyzed nuclei. 3D FISH dramatically shifts the PDF to lower *D* values, while RASER-FISH and CRISPR-Sirius have almost no impact. **D)** Representative images of nuclei labeled with each of the three tested DNA labeling protocols. A chromosome 3 probe was used in the 3D FISH-labeled cell and a chromosome 19 probe was used for the RASER-FISH-labeled cell. The CRISPR-Sirius cell was labeled using a sgRNA targeting the XXYLT1 gene with the MS2 aptamer and stained using Janelia Fluor 646 (JF646). All cells were counterstained with DAPI. The magenta foci are denoted by a circle and an arrow while the nucleus is pseudocolored in blue. Each image is a max projection of ∼15 planes within a z-stack. Scale bar: 5 µm. **E)** Analysis of the labeling efficiency of each tested protocol. Between 15 to 20 confocal images containing nuclei were used to determine the number of foci within each nucleus. The 3D FISH protocol (N=76 nuclei) had the most visible foci, followed by CRISPR-Sirius (N=81 nuclei) had the fewest visible foci, and RASER-FISH (N=99 nuclei). The control sample (N=61 nuclei) had foci detected in one nucleus.

**Table 2.** Comparison of DNA sequence labeling protocol effects on chromatin.

| **Condition** | **Average Nuclear $D \pm$**<br>**Standard Deviation** | **% Change from**<br>**Live Cells** | **% Change from**<br>**Fixed Cells** |
| --- | --- | --- | --- |
| **Live Cells** | 2.56±0.09 (N=104 nuclei) | N/A | N/A |
| **Fixed Cells** | 2.54±0.11 (N=154 nuclei) | -0.79% (p=1) | N/A |
| **3D FISH** | 2.05±0.14<br>(N=168 nuclei) | -19.86%<br>(p=7.42x10<sup>-94</sup>) | -19.22%<br>(p=1.39x10<sup>-108</sup>) |
| **RASER-FISH** | 2.54±0.17<br>(N=74 nuclei) | -0.99%<br>(p=1) | -0.20%<br>(p=1) |
| **CRISPR-Sirius** | 2.48±0.10<br>(N=148 nuclei) | -3.30%<br>(p=1.66x10<sup>-9</sup>) | -2.53 %<br>(p=2.41x10<sup>-6</sup>) |

To determine whether the minimal impact of RASER-FISH and CRISPR-Sirius on average nuclear organization preserves the native chromatin packing domain structure while successfully labeling the targeted foci, we analyzed the histograms of pixel values within nuclei imaged with PWS microscopy and the confocal images of the labeled foci. Like the results presented in Fig 1D, the 3D FISH protocol caused individual nuclei to have a distribution of pixel *D* values centered around an average value of ∼2, although the most common value seen within nuclei was ∼1.74 (Fig 3C). Conversely, the *D* histograms in nuclei labeled with RASER-FISH and CRISPR-Sirius had an average value of ∼2.5 and ∼2.3, with a mode of ∼2.94 and ∼2.64 respectively (Fig 3C). Nuclei labeled with CRISPR-Sirius were also more like live cells qualitatively: there were larger domain structures visible (more high *D* regions that cover a larger portion of the pixels within the nucleus; Fig 3B). While RASER-FISH maintained a similar histogram to CRISPR-Sirius, the nuclei had much smaller domains (Fig 3B), which could be attributed to either the second fixation step used at the end of the protocol or the small volume of formamide contained in the hybridization buffer. To ensure that the minimal impact on chromatin from all three protocols was not due to improper labeling, we performed confocal imaging of cells labeled with each of the three protocols and saw visible foci in several nuclei for each condition (Fig 3D). Interestingly, the DAPI nuclear counterstain showed drastically different chromatin organization. While the 3D FISH labeled cell had an almost uniform fluorescence signal across the nucleus indicating the homogenization of chromatin structure, the RASER-FISH and CRISPR-Sirius labeled cells contained larger variations in DNA interspersed through barren regions (Fig 3D).

Although CRISPR-Sirius had the least significant effect on native chromatin structure and has the added benefit of being a live-cell technique, it is limited to labeling target sequences containing repetitive regions and has a lower labeling efficiency. We used flow cytometry to analyze the percent of cells transduced with CRISPR-Sirius components (i.e., MCP-HALOnls, dCas9, gRNA) and stained with the Janelia Fluor 646 HaloTag ligand. We found that 69% of the transduced population contained a fluorescent signal as compared to 0.3% in the wild-type (WT) control (S1 Fig). We next assessed the impact of the CRISPR-Sirius protocol on chromatin packing domains by imaging a non-targeting empty vector control to compare to WT cells. The empty vector control had little to no impact on the population *D* (WT Δ*D* ≈ –0.02), thus, the transduction and staining steps do not affect nuclear chromatin organization (S2 Fig). To determine whether the length of the DNA target sequence can change the impact of CRISPR-Sirius on chromatin, we tested the effect of using two aptamers (MS2 and PP7) with four single guide RNAs (sgRNAs) for several different pericentromeric and intronic regions, including the pericentromeric region (PR1) of chromosome 19, and repeats on introns of the complement receptor 1 gene (CR1), the xyloside xylosyltransferase gene (XXYLT1), and the fibrillin 3 gene (FBN3). All four sgRNAs caused a similar change in chromatin structure (0.08 < Δ*D* < 0.12) of the same scale as the natural variation that exists within a population (σ*_D_* ≈ 0.08 in the WT control; S2 Fig and S3 Table). Varying the aptamers used with the sgRNA similarly showed trivial differences (S2 Fig and S3 Table), indicating that the choice of aptamer does not affect the changes induced by the protocol on chromatin structure.

The choice of labeling protocol for a study is often based on the relative labeling efficiency, therefore, we analyzed the confocal images taken for all three protocols to determine the proportion of cells with foci. Only one cell in the unstained control had visible foci (2% of the imaged cells), whereas 51% of 3D-FISH labeled cells, 35% of RASER-FISH labeled cells, and 38% of CRISPR-Sirius labeled cells had visible foci (Fig 3E). However, a small number of these cells had three foci (3% of cells in 3D FISH, 1% in RASER-FISH, and 1% in CRISPR-Sirius; Fig 3E). Although all three targets should only have two foci within a single nucleus, the presence of more than two foci could be due to noise in the confocal image, non-specific probe binding, or copy number variation. Taken together, these results show that while RASER-FISH and CRISPR-Sirius are less likely to perturb chromatin structure, they are limited by a reduction in labeling efficiency compared to conventional 3D FISH.

## Discussion

As the reagents and sample preparation procedures in this study use the same steps as other methods for labeling non-chromatin nuclear species (e.g., immunolabeling), knowing how labeling protocols affect the observed structure of the nucleus is of multidisciplinary interest. Every step of the 3D FISH protocol causes significant changes to chromatin in comparison to the live-cell state (Figs 1A-B); this makes developing a protocol that is safe for the study of nanoscale chromatin structure more complex than simply omitting one reagent or step. Furthermore, data obtained from methods such as immunolabeling should be interpreted cautiously. While these labeling techniques typically do not target chromatin, the contents of the entire nucleus are altered by changes to the nuclear environment. We observe widespread changes in chromatin packing in the range of 20-300 nm. Together with the formamide stabilization of the denatured coil phase of DNA, the structural information from 3D FISH on the nanoscale is unlikely to represent the *in vivo* chromatin structure of live cells. We note that the type of imaging modality used could reveal different insights into the effect of 3D FISH on native chromatin structure. In diffraction-limited techniques, such as confocal imaging, it’s possible that the relative spatial positioning of the genome could remain intact [27].

The impact of the 3D FISH protocol is critical for determining whether the locations of genes are altered in super-resolution imaging experiments. If a gene is located within a packing domain, the 3D FISH protocol is unlikely to cause the migration of that gene outside of the domain. However, it does cause the domain to be converted from an actively maintained, heterogeneously structured complex to a swollen, randomly distributed conformation. Solovei et al. showed that sample preparation using the 3D FISH protocol causes the disappearance of heterochromatin, which they hypothesize is becoming more euchromatic [27]. This is consistent with our observed decrease in *D*, and therefore, our results further suggest that this decrease could be ascribed to the disappearance of heterochromatin. Studies that attempt to visualize 3D chromatin structures to compare to results from sequencing-based methodologies could especially be affected by these changes to chromatin domains [10,20,57,58]. For example, techniques such as the optical reconstruction of chromatin architecture (ORCA) [24,25] and Hi-M [23] which aim to probe the internal physical organization of TADs may be impacted by the nanoscale structural artifacts caused by 3D FISH labeling protocols. Although short-range contacts may not be altered, the kilo-base-pair and mega-base-pair interactions are certainly affected by the loss of heterogeneous chromatin structures and heterochromatin.

While we can conclude that the structure of chromatin within the nucleus is dramatically altered by the formamide-based 3D FISH protocol, a limitation of this work is the inability to understand the local structure surrounding an individual fluorescent probe. Additionally, although RASER-FISH removes the need for formamide denaturation to create single-stranded DNA (ssDNA), the use of UV light to induce double-stranded breaks at DNA locations labeled with bromine-nucleotide analogs could change nanoscale chromatin organization at specific loci. This is also a pertinent issue for methods such as CRISPR-Sirius since dCas9 binding results in the bending and unwinding of the target site [59], although we see that the global chromatin structure is not highly altered (Fig 3A). Without ground truth information for the native chromatin organization, it is difficult to determine what changes are induced by the protocol at specific loci. Combining a live cell labeling method with 3D FISH could allow for tracking of foci positioning and changes to surrounding chromatin organization through the protocol. However, our current CRISPR-Sirius labeling methodology has a low signal-to-noise ratio and minimal target fluorescence, requiring confocal imaging with brief periods of laser excitation to prevent photobleaching. An alternative method with low background and high signal stability that allows for repeated imaging of single foci over time, potentially with conventional fluorescence microscopy to reduce the likelihood of photobleaching, would be ideal for this study. Specifically, the SunTag split-sfGFP CRISPR-dCas9 system, which has a low background signal and amplified target fluorescence [60], could be used to label a target sequence in live cells before performing the 3D FISH protocol and imaging with PWS. The ability to use conventional fluorescence microscopy would enable colocalization of fluorescence images with PWS microscopy images for enhanced accuracy in the analysis of alterations to local chromatin structure. Alternatively, as PWS microscopy has limited spatial resolution, a higher-resolution imaging technique that can probe chromatin structure without the use of exogenous labels would be optimal for investigating the local chromatin changes induced by RASER-FISH and CRISPR-Sirius.

Our results offer optimism for alternative DNA labeling techniques such as RASER-FISH and CRISPR/dCas9-based methods. We see that globally, both RASER-FISH and CRISPR-Sirius have minimal impact on chromatin packing domains and the distribution of *D* values within nuclei compared to live, unstained cells. Therefore, while caution is still wise, we do not see severe damage to nanoscale chromatin structure with these techniques, and we hope our findings will encourage a shift from 3D FISH and other formamide-denaturation-based protocols towards RASER-FISH and CRISPR-based approaches.

## Materials and methods

### Cell culture

Human osteosarcoma U2OS cells (ATCC, #HTB-96) were cultured in McCoy’s 5A modified medium (Thermo Fisher Scientific, #16600082) supplemented with 10% fetal bovine serum (FBS; Thermo Fisher Scientific, #26140095) and 100 μg/ml penicillin-streptomycin (Thermo Fisher Scientific, #15140122). HEK293T cells (ATCC, #CRL-1573) were grown in Dulbecco’s Modified Eagle’s Medium (DMEM; Thermo Fisher Scientific, #11965092) supplemented with 10% fetal bovine serum and 100 μg/ml penicillin-streptomycin. All cells were cultured under recommended conditions at 37°C and 5% CO_2_. All cells in this study were maintained between passages 5 and 20. Cells were allowed at least 24::Jhours to re-adhere and recover from trypsin-induced detachment between passages. All cells were negative for mycoplasma contamination (ATCC, #30-1012K) before starting experiments.

### 3D FISH protocol

We followed a modified version of the 3D DNA FISH protocol described by Kocanova, Goiffon, and Bystrcky [29]. Cells were plated in either 12-well glass-bottom plates (Cellvis, #P12-1.5H-N) at 25k seeding density, 6-well glass-bottom plates (Cellvis, #P06-1.5H-N) at 50k, or 35 mm glass-bottom dishes (Cellvis, #D35-14-1-N) at 50k. The plates were then incubated under physiological conditions (5% CO_2_ and 37°C) overnight before use. Cells were quickly rinsed with DPBS (Thermo Fisher Scientific, #14190-144) and then fixed with 4% paraformaldehyde (Electron Microscopy Sciences, #15710) in DPBS for 10 minutes at room temperature. The cells were then washed with DPBS three times for 5 minutes each and treated with freshly made 1 mg/mL sodium borohydride (Fisher Scientific, #S678-10) in DPBS for 10 minutes. The cells were washed with DPBS three times for 5 minutes each, then permeabilized with 0.5% Triton X-100 (Sigma-Aldrich, #93443) in DPBS for 10 minutes at room temperature. Cells were washed three times with DPBS for 5 minutes each and then treated with 0.1 M HCl (Sigma-Aldrich, #320331) diluted with DPBS for 5 minutes at room temperature. After three 5-minute DPBS washes, cells were treated with 0.1 mg/mL RNase A (Sigma-Aldrich, #RNASEA-RO) in DPBS for 45 minutes at 37°C. Cells were then washed with 2x SSC (Sigma-Aldrich, #335266) and incubated in a buffer of 2x SSC, 50% formamide (Sigma-Aldrich, #F7503), and 0.1% Tween 20 (Sigma-Aldrich, #P1379) for 30 minutes at room temperature. Cells were then switched into 10 µL of hybridization buffer composed of 2x SSC, 50% formamide, and 20% dextran sulfate (Sigma-Aldrich, #42867) with 2 µL of a Chromosome 3 Control probe (Empire Genomics, #CHR03-10-RE). Samples were protected from light from hereon. Cells were heated on a heat block at 75°C for 3 minutes, then incubated for between 12-15 hours at 37°C in a humid chamber. After the overnight incubation, cells were washed with a buffer of 2x SSC and 0.1% Tween twice at 60°C for 15 minutes each, and then once at room temperature for 15 minutes. Samples were counterstained using a 10-minute incubation with a 0.1 µg/mL DAPI solution (Thermo Fisher Scientific, #62247) for visualization of nuclei.

### RASER-FISH protocol

We followed the RASER-FISH protocol described by Brown et al [30]. Cells were plated at 25k seeding density in 35 mm glass-bottom dishes. At the time of plating, a 3:1 BrdU/BrdC mix (Sigma-Aldrich, #B5002, and Jena Bioscience, #N-DN-6496) was added to the cells. The plates were then incubated under physiological conditions (5% CO_2_ and 37°C) overnight before use. Cells were washed with DPBS before being fixed in 4% paraformaldehyde and permeabilized in 0.2% Triton X-100 in DPBS for 10 minutes. Cells were incubated with DAPI (1 μg/mL in PBS) and exposed to 254-nm wavelength of ultra-violet (UV) light for 15 minutes, and then treated with Exonuclease III (New England Biolabs, #M0206L) at 37°C for 15 minutes. Dishes were incubated with 2.5 µL of Chromosome 19 Control probe (Empire Genomics, #CHR19-10-RE) in 2.5 µL of *In Situ* Hybridization Buffer (Empire Genomics, #40CJD216) for at least 24 hours at 37°C in a humid chamber. Afterward, cells were washed with 4x SSC, 2x SSC, and then 1x SSC. Cells were then switched to a post-detection fixation solution composed of 16% PFA, 10x DPBS, and Milli-Q water to fix the probe signal. Cells were washed two times with PBST (1x DPBS with 0.05% Tween 20) before staining with a 0.1 µg/mL DAPI solution for 10 minutes and then washed four times with PBST. Finally, cells are rinsed once in DPBS and once in Milli-Q water before being stored in DPBS for imaging.

### CRISPR-Sirius labeling components

Plasmids were obtained from Addgene as bacterial stabs and streaked onto LB-ampicillin plates. After overnight growth and single colony selection, a single colony was inoculated into an LB-ampicillin liquid culture overnight. Plasmid DNA isolation was performed using the QIAprep Spin Miniprep kit (Qiagen, #27104) following the manufacturer’s protocol. pHAGE-TO-dCas9-P2A-HSA (Addgene plasmid #121936), pHAGE-EFS-MCP-HALOnls (Addgene plasmid #121937), pHAGE-EFS-PCP-GFPnls (Addgene plasmid #121938), pLH-sgRNA-Sirius-8XMS2 (Addgene plasmid #121939), pPUR-hU6-sgRNA-Sirius-8XMS2 (Addgene plasmid #121942), and pPUR-mUG-sgRNA-Sirius-8XPP7 (Addgene plasmid #121943) were gifts from Thoru Pederson (UMass Chan Medical School, Worcester, MA) [31]. For labeling targets, one pericentromeric region and three intronic regions were detected. A pericentromeric region (PR1) on the p-arm of chromosome 19 was detected using the target sequence CCnGTTCACTGTCAC (start 21049341, end 21099156, 160 copies) [31] with the forward primer 5’ ACC GGT GAC AGT GAA C 3’ and the reverse primer 5’ AAA CGT TCA CTG TCA C 3’. Repeats on an intronic region of *CR1* encoding for complement receptor 1 (CR1) were detected with the forward primer 5’ ACC GGA GAG GCT GGG 3’ and the reverse primer 5’ AAA CCC CAG CCT CTC C 3’. Xyloside xylosyltransferase 1 (XXYLT1) repeats on an intronic region of the *XXYLT1* gene located on chromosome 3 were detected with the target sequence ATGATATCACAGTGG (start 195199025, end 195233876, 333 copies) [61] with the forward primer 5’ ACC GTG ATA TCA CAG 3’ and the reverse primer 5’ AAA CCT GTG ATA TCA C 3’. Repeats of fibrillin 3 (FBN3) on an intron of the gene *FBN3* located on chromosome 19 were detected with the target sequence ATCCCTCCAACCnGG (start 8201581, end 8202360, 22 copies) [31] with the forward primer 5’ ACC GAT CCC TCC AAC C 3’ and the reverse primer 5’ AAA CGG TTG GAG GGA TC 3’.

### Lentiviral packaging

Lentiviral particles were produced in HEK293T cells using Fugene HD (Promega, #E2311) transfection reagent following the manufacturer’s protocol. The day before transfection, HEK293T cells at low passage (<P10) were split to reach a confluency of 70-80% at the time of transfection. Lentiviral vectors were co-transfected with the lentiviral packaging plasmids pCMV-VSV-G (Addgene plasmid #8454) and pCMV-dR8.2 (Addgene plasmid #8455) (gifts from Robert Weinberg, MIT, Cambridge, MA) [62]. Transfection reactions were assembled in Opti-MEM reduced serum media (Thermo Fisher Scientific, #31985-070). One day before transfection, 1×10^5^ HEK293T cells were plated into each well of a 12-well plate. The following amounts of each plasmid were mixed for packaging each virus: 0.5 μg transfer vector + 0.45 μg pCMV-dR8.2 + 0.05 μg pCMV-VSV-G. The empty vector control was generated by transfecting with the pLH backbone plasmid without a sgRNA target. After 24 hours, the media was changed into fresh DMEM with 10% FBS. Media was refreshed at 12 hours post-transfection, and the virus was harvested at 36-48 hours post-transfection. Viral supernatants were filtered using a 33 mm diameter sterile syringe filter with a 0.45 μm pore size hydrophilic PVDF membrane (Millipore Sigma, #SLHVR33RS) and added to HEK293T cells. Polybrene (8 μg/mL; Sigma-Aldrich, #TR-1003) was supplemented to enhance transduction efficiency. The virus was immediately used or stored at −80°C.

### CRISPR-Sirius transduction

U2OS cells were transduced 6-10 hours after plating. 50k cells were plated in a glass-bottomed 6-well plate. 50 µL dCas9, 50 µL MCP-HALOnls or 50 µL PCP-GFPnls, and 100 µL sgRNA lentiviral particles were added to each well. 24 hours after transduction, lentiviral particles were removed by replacing media. Cells transduced with MCP-HALOnls were incubated for 24 hours before overnight staining with HaloTag-JF646 (Promega, #GA1120) at 10 μM, followed by three washes with DPBS (Thermo Fisher Scientific, #14190-144) and further incubated at 37°C and 5% CO_2_ with phenol-red free media (Cytiva, #SH30270.01) for 24 hours before imaging. Cells were washed three times with DPBS (Gibco, #14190-144) and imaged in phenol-red free media for live cell PWS microscopy. For confocal imaging, cells were fixed with 4% PFA in DPBS for 10 minutes at room temperature and counterstained with 0.1 µg/mL DAPI for 10 minutes after the wash step.

### Confocal microscopy imaging

Confocal images were obtained using the Nikon SoRa Spinning Disk confocal microscope located at the Biological Imaging Facility at Northwestern University in Evanston, IL (RRID: SCR_017767). The Nikon Ti2 inverted microscope is equipped with a Yokogawa CSU-W1 dual-disk spinning disk with 50 µm pinholes, a 50 µm pinhole with micro-lenses SoRa disk, and an ORCA-Fusion Digital CMOS camera (Hamamatsu). Images were collected using a 60x (1.42 NA) oil-immersion objective lens. 3D FISH and RASER-FISH images were obtained using a 2.8x magnifier. For the FISH images, DAPI was excited with a 405 nm laser (50% power with 30 millisecond exposure), and red probes were excited with a 561 nm laser (65% with 200 millisecond exposure). CRISPR-Sirius images were captured with a 1x magnifier. For the CRISPR-Sirius images, DAPI was excited with a 405 nm laser (50% power with 30-millisecond exposure) and HaloTag-JF646 was excited with a 640 nm laser (50% with 200-millisecond exposure). All imaging data were acquired by Nikon NIS Elements acquisition software.

### Confocal microscopy image analysis

To localize the fluorescent puncta, we created a macro using ImageJ software. We first generate max projections of the Z-stack confocal images. To visualize foci, we enhanced the contrast of the images containing the fluorescent probe to saturate 0.001% of the pixels in the image. We then used the DAPI images to generate nuclear masks by first smoothing the images with a Gaussian function with a standard deviation of 10 pixels and then using the auto threshold function with the default parameters. After converting the thresholded image to a mask, we filled holes in the mask and then used the built-in watershed function to split adjacent nuclei. We next filter nuclei by running the analyze particles function to identify objects between 50-1000 pixels in size. To identify foci, we applied a Gaussian function with a 5-pixel standard deviation to smooth the image, found maxima with a prominence of 20, and output a mask with single pixels identifying the center of each maximum. We then counted the number of single points found within each DAPI-defined nuclear mask.

To generate representative confocal images, both the max projection of the confocal images was split into the DAPI and fluorescent probe images. The contrast in both images was enhanced to saturate 0.001% of the pixels within each image. The probe image was then smoothed using a Gaussian function with a standard deviation of 5 pixels. Background subtraction was performed using a rolling ball algorithm with a radius of 5 pixels. Maxima with a prominence of 5000 were identified and all maxima within the tolerance were output as masks. The masks were applied to the contrast-enhanced probe image and then overlaid onto the contrast-enhanced DAPI image.

### PWS microscopy instrumentation

The PWS microscopy optical instrument is built into a Nikon Ti-E inverted microscope with an automated filter turret, a Ti-S-ER sample stage, and a 100x 1.49NA oil objective. Broadband illumination is provided by a white X-Cite 120LED lamp (Excelitas Technologies). To acquire spectral information, a VariSpec liquid crystal tunable filter (LCTF; Cambridge Research & Instrumentation) has been added to the microscope. Spectral data cubes for captured light intensity I(λ, x, y), where λ is the wavelength and (x, y) correspond to pixel positions in the field of view, are collected with a high-speed ORCA-Flash4.0 V3 digital CMOS camera (Hamamatsu) over the wavelength range 500-700 nm at 2 nm intervals [32]. Microscope control and image acquisition are performed using a custom PWS microscopy acquisition plugin (https://github.com/BackmanLab/PWSMicroManager) for µManager [63,64]. For live-cell measurements, cells were imaged under physiological conditions (5% CO_2_ and 37°C) via a stage top incubator (Tokai Hit) with type 37 immersion oil (Electron Microscopy Sciences, #16914-01). Fixed cells were imaged at room temperature (22°C) in trace CO_2_ (open air) conditions using type N immersion oil (Nikon, #MXA22203).

### PWS microscopy imaging and analysis

PWS microscopy measures the spectral standard deviation of internal optical scattering originating from nuclear chromatin, which is related to variations in the refractive index distribution [32,33]. Analysis of spectral data to find the spectral variance is performed using a custom Python code, PWSpy 0.2.13 (https://github.com/BackmanLab/PWSpy), and the associated code for the user interface, pwspy_gui 0.1.13 (https://github.com/BackmanLab/pwspy_gui). Nuclear regions of interest (ROIs) were either hand-drawn for live cells or generated from DAPI images for fixed cells. The calculated spectral variance can be characterized by the mass scaling or chromatin packing scaling, *D*, therefore we can calculate *D* from the variance using a modified autocorrelation function that models chromatin packing domains following power-law scaling within domain boundaries, as described previously [65]. The change in *D* due to each condition is quantified by first averaging *D* within each cell nucleus, and then averaging the average nuclear *D* over hundreds of cells, taken across three biological replicates.

When measuring the effect of individual reagents on changes in *D*, cells were treated with 4% PFA for 10 minutes, washed, and then treated with the individual reagents for the length of time indicated in the 3D FISH protocol. Alternatively, when measuring the effect of the cumulative protocol, the protocol was conducted as indicated, with images being taken after each step (an average of more than three hundred cells taken across three biological replicates). For comparison of fixatives, cells were first imaged live, fixed with the appropriate fixative, imaged fixed, treated with the formamide solution (2x SSC, 50% formamide, and 0.1% Tween 20 for 30 minutes at room temperature), then imaged again. 4% PFA solutions were made in DPBS and added to cells at room temperature for either 10, 30, or 90 minutes. To create the 2% PFA and 2.5% GA (Sigma-Aldrich, #G5882) solution, PFA and GA were mixed in DPBS and added to cells for 10 minutes at room temperature. Both alcohol solutions, 100% methanol (Fisher Scientific, #A452) and 70% ethanol (Sigma-Aldrich, # E7023) in DPBS, were prepared on ice and added to cells for 10 minutes at room temperature. For the optimization of heat and overnight incubation experiment, cells were fixed with 4% PFA for 10 minutes and incubated with formamide solution (2x SSC, 50% formamide, and 0.1% Tween 20 for 30 minutes at room temperature). Cells were then heated on a heating block at either 70°C, 75°C, or 80°C for 3 minutes, then imaged in DPBS. Cells heated at 75°C for 3 minutes were incubated either overnight (between 12-15 hours), for 24 hours, or 48 hours in a humid incubator at 37°C, then imaged in DPBS. In the experiment comparing 3D FISH, RASER-FISH, and CRISPR-Sirius, samples were prepared with the appropriate protocol before imaging with PWS. 3D FISH and RASER-FISH samples underwent fixed cell imaging in DPBS while CRISPR-Sirius samples were imaged live in media. For all experiments, live and 10-minute 4% PFA fixed wild-type cells were used as the control condition (unless otherwise indicated), and three DPBS washes were performed for five minutes each after incubation with each used solution.

To generate representative PWS microscopy images and histograms of *D* pixel intensity within nuclei, each pixel in the PWS microscopy image is first converted to *D* using the methodology described by Eid et al [65]. The nuclear ROIs that were created for each image are applied to the *D* image to extract the nuclear pixels. To plot the histogram of *D* values, all pixels within all nuclei in a condition were combined and plotted. The PWS microscopy images were created by using the nuclear ROIs to pseudocolor nuclei in red and other aspects of the image in gray.

### Flow cytometry

Flow cytometry analysis to determine transduction efficiency was performed on the BD LSRFortessa Cell Analyzer FACSymphony S6 SORP system, located at the Robert H. Lurie Comprehensive Cancer Center Flow Cytometry Core Facility at Northwestern University in Evanston, IL. For all FACS analyses the same protocol was used. 48 hours post-transduction, cells were harvested and fixed. Briefly, cells were washed with DPBS, trypsinized (Gibco, #25200-056), neutralized with media, and then centrifuged at 500 x g for 5 minutes. Cells were then resuspended in 500 μL of 2% PFA and DPBS and fixed for 10 minutes at room temperature, followed by centrifugation and resuspension in cold FACS buffer composed of DPBS with 1% of BSA and 2mM EDTA (Thermo Fisher Scientific, #1860851) at 4°C until analysis could be performed the following day. At least 10, 000 events were recorded. Flow cytometric data were analyzed using FlowJo software version 10.6.1. Gating to identify fluorescent cells was performed using negative controls, and then applied across all collected samples.

### Quantification and statistical analysis

Statistical analysis was performed in Python 3.9.13 using the SciPy 1.9.3 [66] and statsmodels 0.14.1 packages. We used the multiple comparison function to run pairwise t-tests on all sample pairs with a Bonferroni correction. Data are representative of at least three independent experiments, each of which contained at least one hundred cells per treatment. All data are presented as the mean ± standard deviation. Bonferroni corrected P values are reported. A p-value of <0.05 was considered statistically significant.

## Author contributions

A.R.S. conceived the project. A.R.S., J.F., and I.C.Y. performed the 3D FISH experiments. T.K. performed the RASER-FISH experiments. S.J. designed and assisted in the implementation of CRISPR-Sirius labeling. A.R.S. and E.M.P. performed the CRISPR-Sirius experiments. N.M.A. developed the PWS microscopy imaging and analysis software. A.R.S., J.F., E.M.P., I.C.Y., and J.A.P. performed PWS microscopy imaging. A.R.S., J.F., E.M.P., T.K., and I.C.Y. analyzed PWS microscopy images. E.M.P., T.K., I.C.Y., C.D., and P.C.G. performed confocal imaging. E.M.P. performed flow cytometry. J.F. analyzed confocal images and flow cytometry data. A.R.S., J.F., and E.M.P. prepared the manuscript. All authors edited the manuscript. V.B. and I.S. supervised the project.

## Acknowledgments

We thank David VanDerway for aiding in culturing U2OS cells. We thank the Robert H. Lurie Comprehensive Cancer Center of Northwestern University in Chicago, IL, for the use of the Flow Cytometry Core Facility, which provided flow cytometric analysis training and use of the BD LSRFortessa Cell Analyzer FACSymphony S6 SORP system. We thank the Biological Imaging Facility at Northwestern University (RRID: SCR_017767) for providing training and use of the Nikon SoRa Spinning Disc confocal microscope. We thank Dr. Thoru Pederson for sharing CRISPR-Sirius plasmids with us and Dr. Robert Weinberg for sharing the lentiviral packaging plasmids. This work was supported by funding from the National Science Foundation under the NSF EFRI (EFMA-1830961 awarded to PI V.B. and co-PI I.S.), the National Institutes of Health through the National Cancer Institute (R01CA228272 awarded to V.B. and I.S., R01CA225002 awarded to V.B., U54268084 awarded to PI V.B. and co-PI I.S., and U54CA261964 awarded to core lead V.B.), the Center for Physical Genomics and Engineering at Northwestern University, and philanthropic support from K. Hudson and R. Goldman, S. Brice and J. Esteve, M. E. Holliday and I. Schneider, the Christina Carinato Charitable Foundation, and D. Sachs. The Biological Imaging Facility at Northwestern University is graciously supported by the Chemistry for Life Processes Institute, the NU Office for Research, the Department of Molecular Biosciences, and the Rice Foundation. The Lurie Cancer Center is supported in part by an NCI Cancer Center Support Grant #P30 CA060553. This material is based upon work performed by J.F. which was supported by the National Science Foundation Graduate Research Fellowship under Grant No. DGE-1842165. Any opinions, findings, conclusions, or recommendations expressed in this material are those of the authors and do not necessarily reflect the views of the National Science Foundation. The funders had no role in the study design, data collection, data analysis, the decision to publish, or the preparation of the manuscript.

## Supporting information

**S1. Table.**
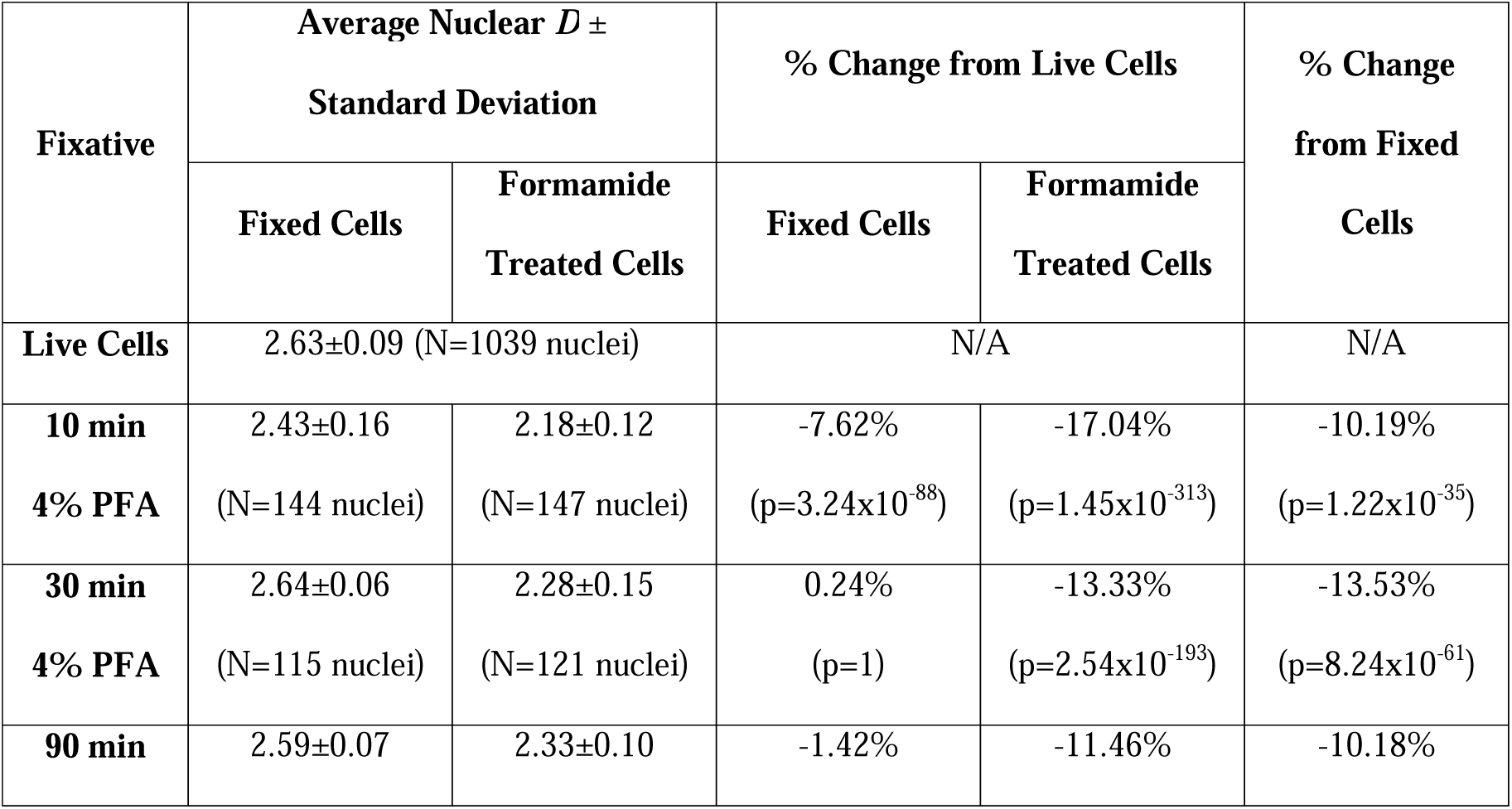
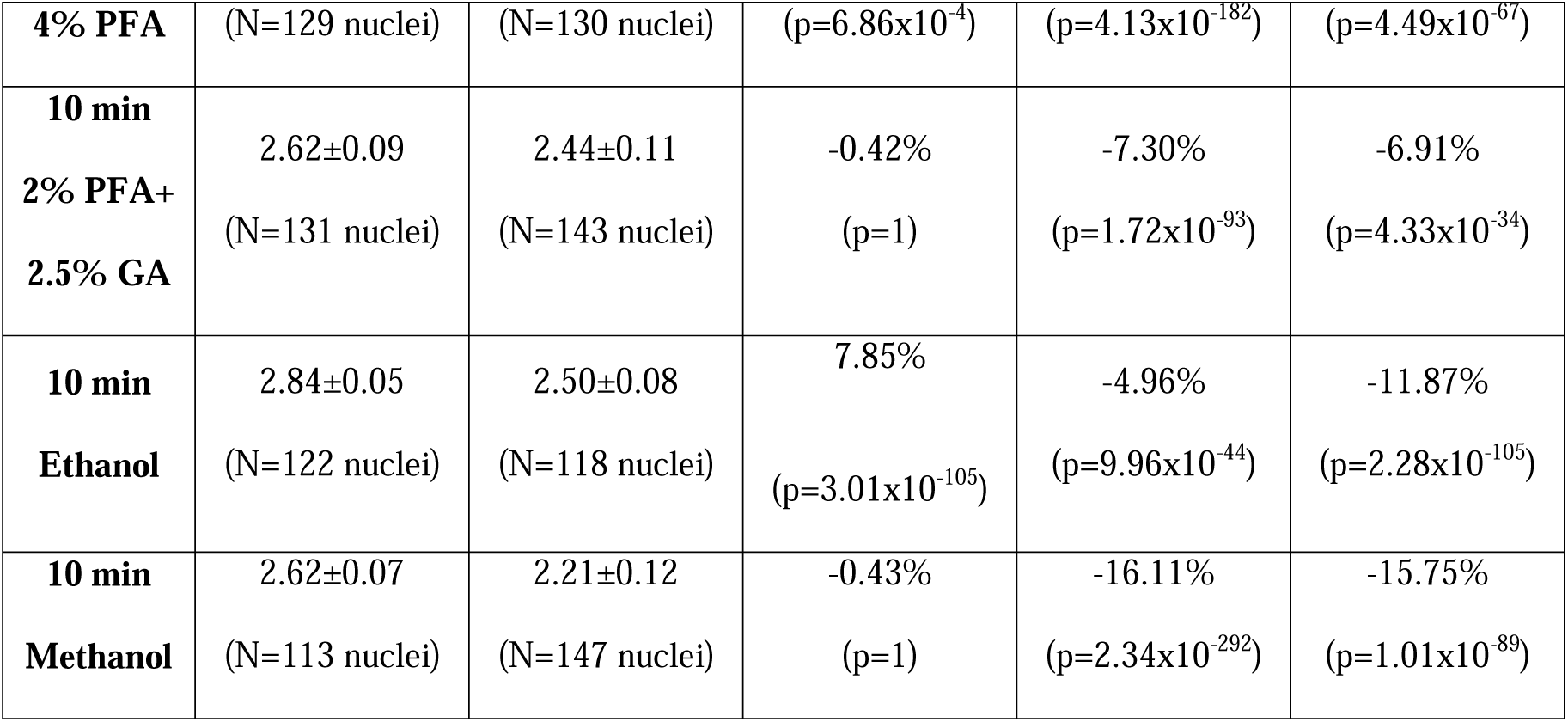
Effect of formamide on cells fixed with different reagents and incubation lengths.

**S2. Table.**
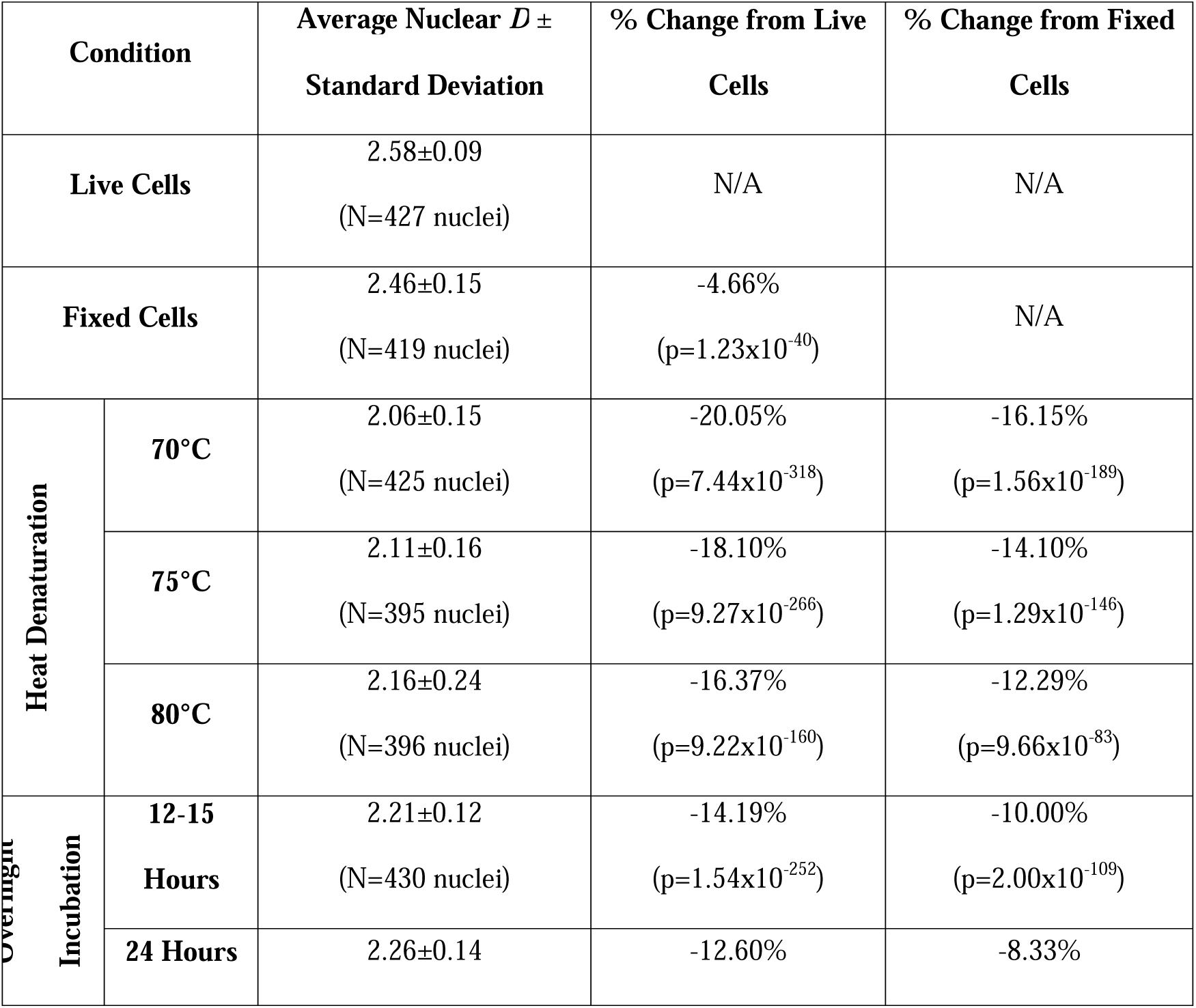
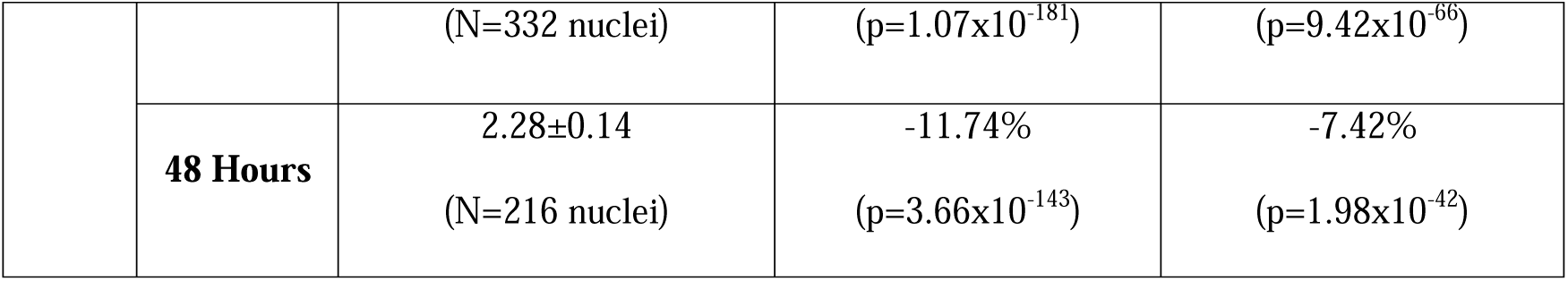
Optimization of the heat and overnight steps do not fully recover the original chromatin structures.

**S3. Table.**
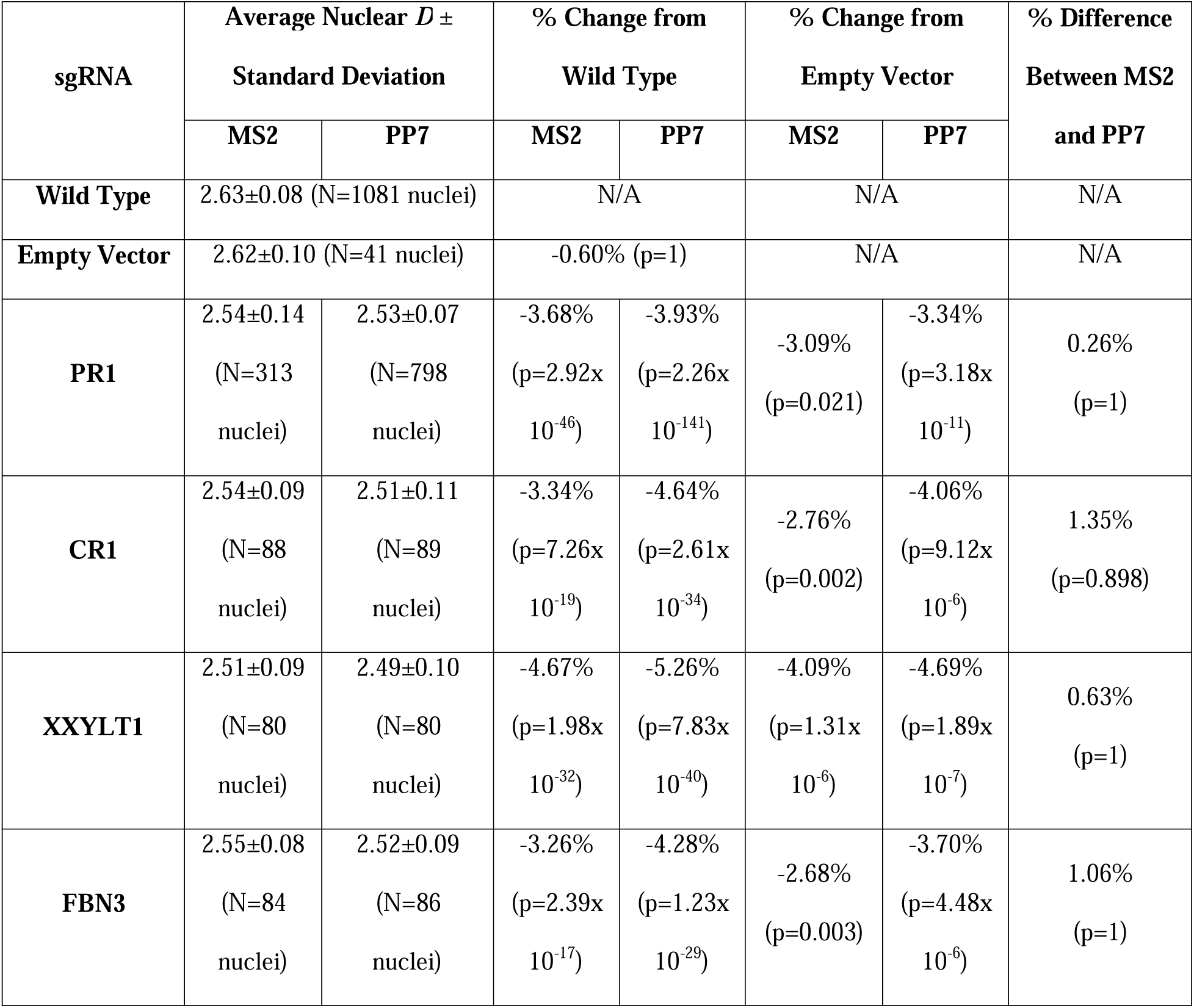
Effect of sgRNA and aptamer on chromatin structure.

**S1. Fig.**
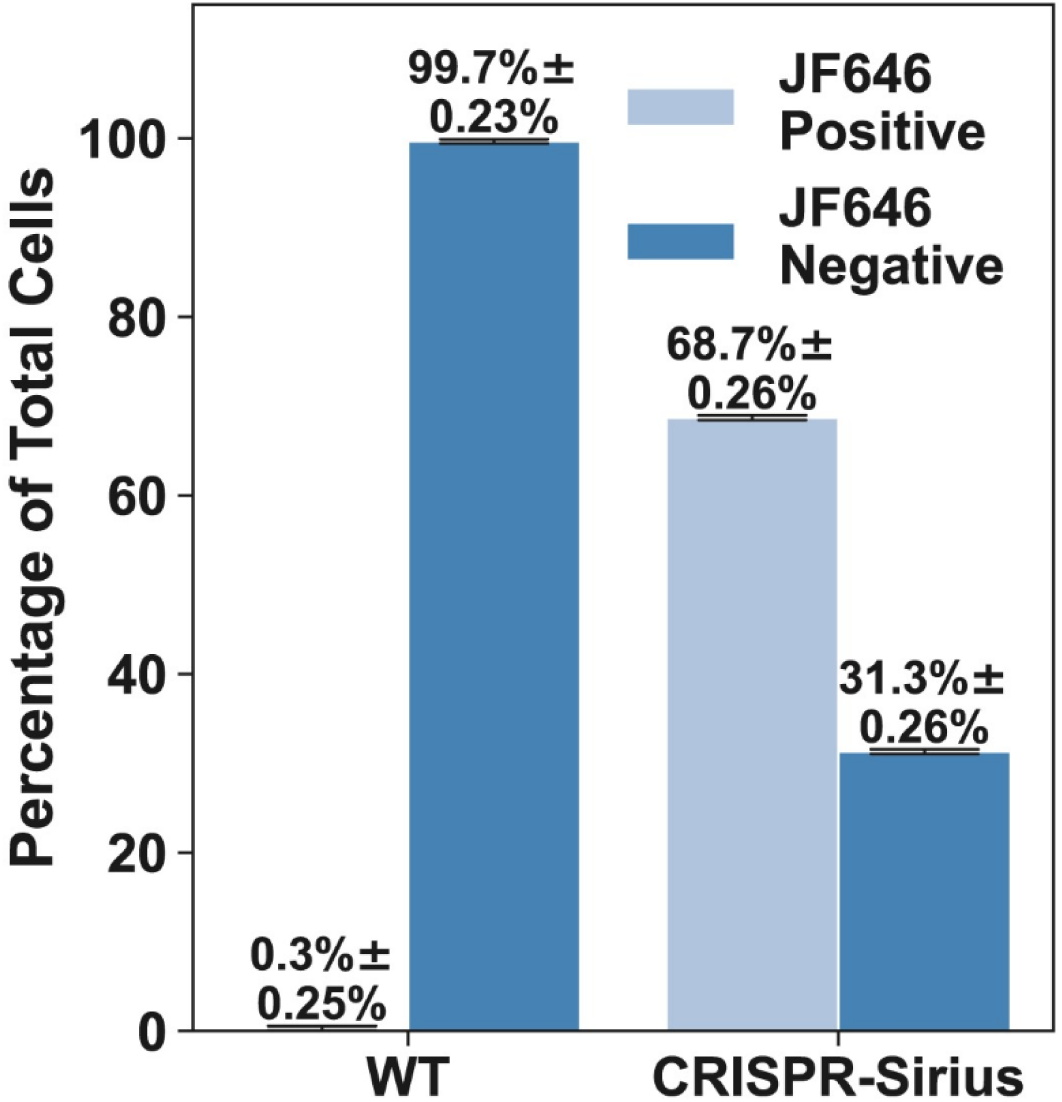
Transduction efficiency of CRISPR-Sirius. CRISPR-Sirius transduction efficiency as determined by the number of cells exhibiting fluorescence measured using flow cytometry. The XXYLT1 gene was labeled with the MS2 aptamer and stained using Janelia Fluorophore 646 for visualization. In the control sample, ∼80,000 live cells were captured by flow cytometry, while the CRISPR-Sirius sample had ∼70,000 live cells.

**S2. Fig.**
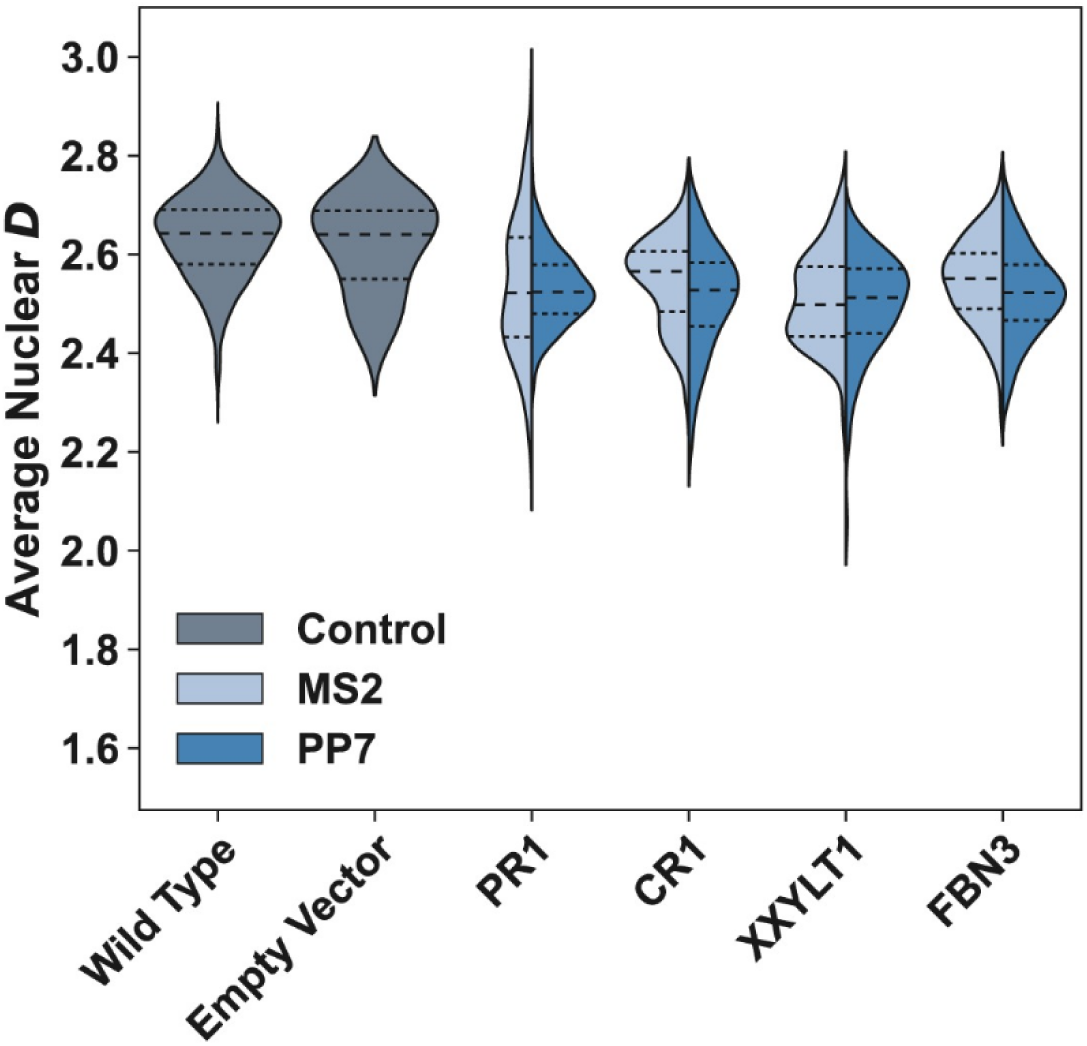
Impact of the CRISPR-Sirius aptamer and sgRNA on chromatin structure. PWS microscopy imaging of cells labeled with CRISPR-Sirius using two different aptamers (MS2 and PP7) and four primers targeting the pericentromeric region (PR1) on chromosome 19, repeats on the gene which codes for complement receptor 1 (CR1) on chromosome 1, an intronic region on the gene for xyloside xylosyltransferase 1 (XXYLT1) on chromosome 3, and repeats on an intron for the gene encoding fibrillin 3 (FBN3) on chromosome 19. Average nuclear *D* decreases slightly for both aptamers and all primers, however, all CRISPR populations reach similar mean *D* values after lentiviral transduction, showing that the choice of guide RNA and aptamer is not the driving factor for chromatin changes. Violins include data from between 40-1100 nuclei. Dashed lines within violins denote the 75^th^ percentile, median, and 25^th^ percentile from top to bottom.

## References

1. Zhao H, Zhang Y, Zhang SB, Jiang C, He QY, Li MQ, et al. The structure of the nucleosome core particle of chromatin in chicken erythrocytes visualized by using atomic force microscopy. Cell Res. 1999;9: 255–260. doi:10.1038/sj.cr.7290024

2. Dixon JR, Selvaraj S, Yue F, Kim A, Li Y, Shen Y, et al. Topological domains in mammalian genomes identified by analysis of chromatin interactions. Nature. 2012;485: 376–380. doi:10.1038/nature11082

3. Dali R, Blanchette M. A critical assessment of topologically associating domain prediction tools. Nucleic Acids Res. 2017;45: 2994–3005. doi:10.1093/nar/gkx145

4. Lieberman-Aiden E, van Berkum NL, Williams L, Imakaev M, Ragoczy T, Telling A, et al. Comprehensive Mapping of Long-Range Interactions Reveals Folding Principles of the Human Genome. Science. 2009;326: 289–293. doi:10.1126/science.1181369

5. Quinodoz SA, Ollikainen N, Tabak B, Palla A, Schmidt JM, Detmar E, et al. Higher-Order Inter-chromosomal Hubs Shape 3D Genome Organization in the Nucleus. Cell. 2018;174: 744–757.e24. doi:10.1016/j.cell.2018.05.024

6. Shanta O, Noor A, Chaisson MJP, Sanders AD, Zhao X, Malhotra A, et al. The effects of common structural variants on 3D chromatin structure. BMC Genomics. 2020;21: 95. doi:10.1186/s12864-020-6516-1

7. Miron E, Oldenkamp R, Brown JM, Pinto DMS, Xu CS, Faria AR, et al. Chromatin arranges in chains of mesoscale domains with nanoscale functional topography independent of cohesin. Sci Adv. 2020;6: eaba8811. doi:10.1126/sciadv.aba8811

8. Li Y, Eshein A, Virk RKA, Eid A, Wu W, Frederick J, et al. Nanoscale chromatin imaging and analysis platform bridges 4D chromatin organization with molecular function. Sci Adv. 2021;7: eabe4310. doi:10.1126/sciadv.abe4310

9. Heo S-J, Thakur S, Chen X, Loebel C, Xia B, McBeath R, et al. Aberrant chromatin reorganization in cells from diseased fibrous connective tissue in response to altered chemomechanical cues. Nat Biomed Eng. 2023;7: 177–191. doi:10.1038/s41551-022-00910-5

10. Bintu B, Mateo LJ, Su J-H, Sinnott-Armstrong NA, Parker M, Kinrot S, et al. Super-resolution chromatin tracing reveals domains and cooperative interactions in single cells. Science. 2018;362: eaau1783. doi:10.1126/science.aau1783

11. Li Y, Agrawal V, Virk RKA, Roth E, Li WS, Eshein A, et al. Analysis of three-dimensional chromatin packing domains by chromatin scanning transmission electron microscopy (ChromSTEM). Sci Rep. 2022;12: 12198. doi:10.1038/s41598-022-16028-2

12. Almassalha LM, Bauer GM, Wu W, Cherkezyan L, Zhang D, Kendra A, et al. Macrogenomic engineering via modulation of the scaling of chromatin packing density. Nat Biomed Eng. 2017;1: 902–913. doi:10.1038/s41551-017-0153-2

13. Almassalha LM, Tiwari A, Ruhoff PT, Stypula-Cyrus Y, Cherkezyan L, Matsuda H, et al. The Global Relationship between Chromatin Physical Topology, Fractal Structure, and Gene Expression. Sci Rep. 2017;7: 41061. doi:10.1038/srep41061

14. Virk RKA, Wu W, Almassalha LM, Bauer GM, Li Y, VanDerway D, et al. Disordered chromatin packing regulates phenotypic plasticity. Sci Adv. 2020;6: eaax6232. doi:10.1126/sciadv.aax6232

15. Nielsen B, Albregtsen F, Danielsen HE. The Use of Fractal Features from the Periphery of Cell Nuclei as a Classification Tool. Anal Cell Pathol. 1999;19: 21–37. doi:10.1155/1999/986086

16. Stypula-Cyrus Y, Damania D, Kunte DP, Cruz MD, Subramanian H, Roy HK, et al. HDAC Up-Regulation in Early Colon Field Carcinogenesis Is Involved in Cell Tumorigenicity through Regulation of Chromatin Structure. PLOS ONE. 2013;8: e64600. doi:10.1371/journal.pone.0064600

17. Bauer GM, Stypula-Cyrus Y, Subramanian H, Cherkezyan L, Viswanathan P, Zhang D, et al. The transformation of the nuclear nanoarchitecture in human field carcinogenesis. Future Sci OA. 2017;3: FSO206. doi:10.4155/fsoa-2017-0027

18. Goutzanis L, Papadogeorgakis N, Pavlopoulos PM, Katti K, Petsinis V, Plochoras I, et al. Nuclear fractal dimension as a prognostic factor in oral squamous cell carcinoma. Oral Oncol. 2008;44: 345–353. doi:10.1016/j.oraloncology.2007.04.005

19. Metze K. Fractal dimension of chromatin: potential molecular diagnostic applications for cancer prognosis. Expert Rev Mol Diagn. 2013;13: 719–735. doi:10.1586/14737159.2013.828889

20. Boettiger AN, Bintu B, Moffitt JR, Wang S, Beliveau BJ, Fudenberg G, et al. Super-resolution imaging reveals distinct chromatin folding for different epigenetic states. Nature. 2016;529: 418–422. doi:10.1038/nature16496

21. Ou HD, Phan S, Deerinck TJ, Thor A, Ellisman MH, O’Shea CC. ChromEMT: Visualizing 3D chromatin structure and compaction in interphase and mitotic cells. Science. 2017;357: eaag0025. doi:10.1126/science.aag0025

22. Beliveau BJ, Boettiger AN, Nir G, Bintu B, Yin P, Zhuang X, et al. In Situ Super-Resolution Imaging of Genomic DNA with OligoSTORM and OligoDNA-PAINT. In: Erfle H, editor. Super-Resolution Microscopy: Methods and Protocols. New York, NY: Springer; 2017. pp. 231–252. doi:10.1007/978-1-4939-7265-4_19

23. Cardozo Gizzi AM, Cattoni DI, Fiche J-B, Espinola SM, Gurgo J, Messina O, et al. Microscopy-Based Chromosome Conformation Capture Enables Simultaneous Visualization of Genome Organization and Transcription in Intact Organisms. Mol Cell. 2019;74: 212–222.e5. doi:10.1016/j.molcel.2019.01.011

24. Mateo LJ, Murphy SE, Hafner A, Cinquini IS, Walker CA, Boettiger AN. Visualizing DNA folding and RNA in embryos at single-cell resolution. Nature. 2019;568: 49–54. doi:10.1038/s41586-019-1035-4

25. Mateo LJ, Sinnott-Armstrong N, Boettiger AN. Tracing DNA paths and RNA profiles in cultured cells and tissues with ORCA. Nat Protoc. 2021;16: 1647–1713. doi:10.1038/s41596-020-00478-x

26. Mongelard F, Vourc’h C, Robert-Nicoud M, Usson Y. Quantitative assessment of the alteration of chromatin during the course of FISH procedures. Cytometry. 1999;36: 96–101.

27. Solovei I, Cavallo A, Schermelleh L, Jaunin F, Scasselati C, Cmarko D, et al. Spatial Preservation of Nuclear Chromatin Architecture during Three-Dimensional Fluorescence in Situ Hybridization (3D-FISH). Exp Cell Res. 2002;276: 10–23. doi:10.1006/excr.2002.5513

28. Markaki Y, Smeets D, Fiedler S, Schmid VJ, Schermelleh L, Cremer T, et al. The potential of 3D-FISH and super-resolution structured illumination microscopy for studies of 3D nuclear architecture. BioEssays. 2012;34: 412–426. doi:10.1002/bies.201100176

29. Kocanova S, Goiffon I, Bystricky K. 3D FISH to analyse gene domain-specific chromatin re-modeling in human cancer cell lines. Methods. 2018;142: 3–15. doi:10.1016/j.ymeth.2018.02.013

30. Brown JM, De Ornellas S, Parisi E, Schermelleh L, Buckle VJ. RASER-FISH: non-denaturing fluorescence in situ hybridization for preservation of three-dimensional interphase chromatin structure. Nat Protoc. 2022;17: 1306–1331. doi:10.1038/s41596-022-00685-8

31. Ma H, Tu L-C, Naseri A, Chung Y-C, Grunwald D, Zhang S, et al. CRISPR-Sirius: RNA scaffolds for signal amplification in genome imaging. Nat Methods. 2018;15: 928–931. doi:10.1038/s41592-018-0174-0

32. Almassalha LM, Bauer GM, Chandler JE, Gladstein S, Cherkezyan L, Stypula-Cyrus Y, et al. Label-free imaging of the native, living cellular nanoarchitecture using partial-wave spectroscopic microscopy. Proc Natl Acad Sci. 2016;113: E6372–E6381. doi:10.1073/pnas.1608198113

33. Cherkezyan L, Subramanian H, Backman V. What structural length scales can be detected by the spectral variance of a microscope image? Opt Lett. 2014;39: 4290– 4293. doi:10.1364/OL.39.004290

34. Gennes P-G de. Scaling Concepts in Polymer Physics. Cornell University Press; 1979.

35. Flory PJ. The Configuration of Real Polymer Chains. J Chem Phys. 1949;17: 303–310. doi:10.1063/1.1747243

36. Sanchez IC. Phase Transition Behavior of the Isolated Polymer Chain. Macromolecules. 1979;12: 980–988. doi:10.1021/ma60071a040

37. Yokota H, van den Engh G, Hearst JE, Sachs RK, Trask BJ. Evidence for the organization of chromatin in megabase pair-sized loops arranged along a random walk path in the human G0/G1 interphase nucleus. J Cell Biol. 1995;130: 1239– 1249. doi:10.1083/jcb.130.6.1239

38. Sachs RK, van den Engh G, Trask B, Yokota H, Hearst JE. A random-walk/giant-loop model for interphase chromosomes. Proc Natl Acad Sci. 1995;92: 2710–2714. doi:10.1073/pnas.92.7.2710

39. Huang K, Li Y, Shim AR, Virk RKA, Agrawal V, Eshein A, et al. Physical and data structure of 3D genome. Sci Adv. 2020;6: eaay4055. doi:10.1126/sciadv.aay4055

40. Shim AR, Huang K, Backman V, Szleifer I. Chromatin as self-returning walks: From population to single cell and back. Biophys Rep. 2022;2: 100042. doi:10.1016/j.bpr.2021.100042

41. Lebedev D v., Filatov M v., Kuklin A i., Islamov AKh, Kentzinger E, Pantina R, et al. Fractal nature of chromatin organization in interphase chicken erythrocyte nuclei: DNA structure exhibits biphasic fractal properties. FEBS Lett. 2005;579: 1465–1468. doi:10.1016/j.febslet.2005.01.052

42. Bancaud A, Lavelle C, Huet S, Ellenberg J. A fractal model for nuclear organization: current evidence and biological implications. Nucleic Acids Res. 2012;40: 8783–8792. doi:10.1093/nar/gks586

43. Bancaud A, Huet S, Daigle N, Mozziconacci J, Beaudouin J, Ellenberg J. Molecular crowding affects diffusion and binding of nuclear proteins in heterochromatin and reveals the fractal organization of chromatin. EMBO J. 2009;28: 3785–3798. doi:10.1038/emboj.2009.340

44. Li Y, Almassalha LM, Chandler JE, Zhou X, Stypula-Cyrus YE, Hujsak KA, et al. The effects of chemical fixation on the cellular nanostructure. Exp Cell Res. 2017;358: 253–259. doi:10.1016/j.yexcr.2017.06.022

45. Zhou X, Gladstein S, Almassalha LM, Li Y, Eshein A, Cherkezyan L, et al. Preservation of cellular nano-architecture by the process of chemical fixation for nanopathology. PLOS ONE. 2019;14: e0219006. doi:10.1371/journal.pone.0219006

46. Hepperger C, Otten S, von Hase J, Dietzel S. Preservation of large-scale chromatin structure in FISH experiments. Chromosoma. 2007;116: 117–133. doi:10.1007/s00412-006-0084-2

47. Cifra P, Karasz FE, MacKnight WJ. Expansion of polymer coils in miscible polymer blends of asymmetric composition. Macromolecules. 1992;25: 192–194. doi:10.1021/ma00027a032

48. Hammouda B, Worcester D. The Denaturation Transition of DNA in Mixed Solvents. Biophys J. 2006;91: 2237–2242. doi:10.1529/biophysj.106.083691

49. Blake RD, Delcourt SG. Thermodynamic effects of formamide on DNA stability. Nucleic Acids Res. 1996;24: 2095–2103.

50. Finn EH, Misteli T. A high-throughput DNA FISH protocol to visualize genome regions in human cells. STAR Protoc. 2021;2: 100741. doi:10.1016/j.xpro.2021.100741

51. Miething F, Hering S, Hanschke B, Dressler J. Effect of Fixation to the Degradation of Nuclear and Mitochondrial DNA in Different Tissues. J Histochem Cytochem. 2006;54: 371–374. doi:10.1369/jhc.5B6726.2005

52. Falconi M, Teti G, Zago M, Pelotti S, Gobbi P, Breschi L, et al. Effect of Fixative on Chromatin Structure and DNA Detection. Microsc Res Tech. 2007;70: 599–606. doi:10.1002/jemt.20440

53. Hobro AJ, Smith NI. An evaluation of fixation methods: Spatial and compositional cellular changes observed by Raman imaging. Vib Spectrosc. 2017;91: 31–45. doi:10.1016/j.vibspec.2016.10.012

54. Gillespie JW, Best CJM, Bichsel VE, Cole KA, Greenhut SF, Hewitt SM, et al. Evaluation of Non-Formalin Tissue Fixation for Molecular Profiling Studies. Am J Pathol. 2002;160: 449–457. doi:10.1016/S0002-9440(10)64864-X

55. McConaughy BL, Laird CD, McCarthy BJ. Nucleic acid reassociation in formamide. Biochemistry. 1969;8: 3289–3295. doi:10.1021/bi00836a024

56. Brown JM, Roberts NA, Graham B, Waithe D, Lagerholm C, Telenius JM, et al. A tissue-specific self-interacting chromatin domain forms independently of enhancer-promoter interactions. Nat Commun. 2018;9: 3849. doi:10.1038/s41467-018-06248-4

57. Giorgetti L, Galupa R, Nora EP, Piolot T, Lam F, Dekker J, et al. Predictive Polymer Modeling Reveals Coupled Fluctuations in Chromosome Conformation and Transcription. Cell. 2014;157: 950–963. doi:10.1016/j.cell.2014.03.025

58. Stevens TJ, Lando D, Basu S, Atkinson LP, Cao Y, Lee SF, et al. 3D structures of individual mammalian genomes studied by single-cell Hi-C. Nature. 2017;544: 59–64. doi:10.1038/nature21429

59. Yamano T, Nishimasu H, Zetsche B, Hirano H, Slaymaker IM, Li Y, et al. Crystal Structure of Cpf1 in Complex with Guide RNA and Target DNA. Cell. 2016;165: 949–962. doi:10.1016/j.cell.2016.04.003

60. Chaudhary N, Nho S-H, Cho H, Gantumur N, Ra JS, Myung K, et al. Background-suppressed live visualization of genomic loci with an improved CRISPR system based on a split fluorophore. Genome Res. 2020;30: 1306–1316. doi:10.1101/gr.260018.119

61. Pujadas EM, Wei X, Acosta N, Carter L, Yang J, Almassalha L, et al. Depletion of lamins B1 and B2 alters chromatin mobility and induces differential gene expression by a mesoscale-motion dependent mechanism. bioRxiv; 2023. p. 2023.06.26.546573. doi:10.1101/2023.06.26.546573

62. Stewart SA, Dykxhoorn DM, Palliser D, Mizuno H, Yu EY, An DS, et al. Lentivirus-delivered stable gene silencing by RNAi in primary cells. RNA. 2003;9: 493–501. doi:10.1261/rna.2192803

63. Edelstein A, Amodaj N, Hoover K, Vale R, Stuurman N. Computer Control of Microscopes Using µManager. Curr Protoc Mol Biol. 2010;92: 14.20.1–14.20.17. doi:10.1002/0471142727.mb1420s92

64. Edelstein AD, Tsuchida MA, Amodaj N, Pinkard H, Vale RD, Stuurman N. Advanced methods of microscope control using μManager software. J Biol Methods. 2014;1: e10. doi:10.14440/jbm.2014.36

65. Eid A, Eshein A, Li Y, Virk R, Derway DV, Zhang D, et al. Characterizing chromatin packing scaling in whole nuclei using interferometric microscopy. Opt Lett. 2020;45: 4810–4813. doi:10.1364/OL.400231

66. Virtanen P, Gommers R, Oliphant TE, Haberland M, Reddy T, Cournapeau D, et al. SciPy 1.0: fundamental algorithms for scientific computing in Python. Nat Methods. 2020;17: 261–272. doi:10.1038/s41592-019-0686-2

